# Zika virus remodels and hijacks IGF2BP2 ribonucleoprotein complex to promote viral replication organelle biogenesis

**DOI:** 10.1101/2023.12.08.570783

**Authors:** Clément Mazeaud, Stefan Pfister, Jonathan E. Owen, Higor Sette Pereira, Flavie Charbonneau, Zachary E. Robinson, Anaïs Anton, Cheyanne L. Bemis, Aïssatou Aïcha Sow, Trushar R. Patel, Christopher J. Neufeldt, Pietro Scaturro, Laurent Chatel-Chaix

## Abstract

Zika virus (ZIKV) infection causes significant human disease that, with no approved treatment or vaccine, constitutes a major public health concern. Its life cycle entirely relies on the cytoplasmic fate of the viral RNA genome (vRNA) through a fine-tuned equilibrium between vRNA translation, replication and packaging into new virions, all within virus-induced replication organelles (vRO). In this study, with an RNAi mini-screening and subsequent functional characterization, we have identified insulin-like growth factor 2 mRNA-binding protein 2 (IGF2BP2) as a new host dependency factor that regulates vRNA synthesis. In infected cells, IGF2BP2 associates with viral NS5 polymerase and redistributes to the perinuclear viral replication compartment. Combined fluorescence *in situ* hybridization-based confocal imaging, *in vitro* binding assays, and immunoprecipitation coupled to RT-qPCR, showed that IGF2BP2 directly interacts with ZIKV vRNA 3’-nontranslated region. Using ZIKV sub-genomic replicons and a replication-independent vRO induction system, we demonstrated that IGF2BP2 knockdown impairs *de novo* viral organelle biogenesis and, consistently, vRNA synthesis. Finally, the analysis of immunopurified IGF2BP2 complex using quantitative mass spectrometry and RT-qPCR, revealed that ZIKV infection alters the protein and RNA interactomes of IGF2BP2. Altogether, our data support that ZIKV hijacks and remodels the IGF2BP2 ribonucleoprotein complex to regulate vRO biogenesis and vRNA neosynthesis.

## INTRODUCTION

Zika virus (ZIKV) is the causative agent in several epidemics in the last 10 years, with the biggest outbreak in Latin America in 2015, making ZIKV a major public health concern. ZIKV belongs to the *Orthoflavivirus* genus within the *Flaviviridae* family which comprises over 70 arthropod-borne viruses (arboviruses) such as dengue virus (DENV), West Nile virus (WNV) and yellow fever virus. Similar to the closely related DENV, ZIKV is primarily transmitted to humans by *Aedes* spp mosquito bites (1) and can cause Guillain-Barré syndrome in adults. When vertically transmitted to a fetus during pregnancy, this neurotropic virus can infect the placenta as well as neural progenitor cells, microglial cells and astrocytes in the developing brain, which may eventually cause severe microcephaly in newborns (2–7). Unfortunately, no antiviral treatments against ZIKV or prophylactic vaccines are currently approved. It is thus important to better understand the molecular mechanisms controlling the ZIKV replication in order to identify new therapeutic targets. ZIKV is a single-stranded positive sense enveloped RNA virus. Its ∼11 kb-long viral RNA (vRNA) genome is composed of an open reading frame flanked by 5’ and 3’nontranslated regions (NTR). The vRNA plays a central role in different steps of the viral replication cycle including being translated to produce all viral proteins, is used as template for genome neosynthesis and is selectively packaged into assembling viral particles. The vRNA, translated at the ER membrane, encodes one polyprotein which is matured into 10 viral proteins. Three structural proteins (C, prM and E), together with vRNA assemble new viral particles, and seven nonstructural proteins (NS1, NS2A, NS2B, NS3, NS4A, NS4B and NS5) are responsible for vRNA neosynthesis, via the activity of the RNA-dependent RNA polymerase (RdRp) NS5 along with the contribution of other viral proteins and host factors (8,9). To efficiently replicate, orthoflaviviruses have evolved to establish a fine-tuned equilibrium between vRNA translation, replication and encapsidation steps since they cannot occur simultaneously. However, the mechanisms underlying the spatio-temporal fate of vRNA remain elusive. These processes occur in viral replication organelles (vRO) which result from major alterations of the endoplasmic reticulum (ER) and are believed to coordinate in time and space the steps of the life cycle, notably through the spatial segregation of the vRNA, at different steps of the life cycle. vROs accumulate in a large cage-like compartment which is located in the perinuclear area (10). Three architectures of these viral organelles-like ultrastructures have been described: vesicle packets (VP), virus bags (VB), and convoluted membranes (CM). The VPs resulting from ER invagination, are spherical vesicles with a diameter of approximately 90 nm connected to the cytosol through a 10 nm-wide pore-like opening (10–13). They are believed to host vRNA synthesis. Indeed, the non-structural proteins NS5, NS3, NS1, NS4A and NS4B, absolutely required for vRNA replication, and also double-stranded RNA (dsRNA), the viral RNA replication intermediate, are enriched in VPs as imaged by immunogold labeling followed by transmission electron microscopy (11,14). It has been proposed that the RNA genome exit VPs through their pore to be directly packaged into assembling particles budding into the ER lumen. Immature virions geometrically accumulate in dilated ER cisternae referred to as virus bags (VB). The viral and cellular determinants of vRO biogenesis are poorly understood mostly because this process will be impacted (even indirectly) by any perturbations of viral replication efficiency and hence, viral factors abundance. The recent advent of replication-independent VP induction systems identified several determinants of *de novo* VP biogenesis, most notably including, ZIKV NS4A, NS1 and 3’NTR RNA, as well as host factors ATL2, VCP and RACK1 (15–21).

Multiple reports have shown that cellular RNA-binding proteins (RBP) regulate the replication, translation and/or encapsidation of vRNA often through associations with 5’ or 3’ NTRs (for review, see (8)). For instance, YBX1 associates with 3’ NTR of DENV vRNA and regulates both vRNA translation and viral particle production (22–24). However, it remains unclear whether all these proteins are co-opted and regulate replication within a single ribonucleprotein (RNP) complex. In case of the hepatitis C virus (HCV), a *Flaviviridae* from the *Hepacivirus* genus, it was shown that the YBX1 RNP comprising HCV vRNA, IGF2BP2, LARP1, DDX6 and other cell proteins regulates the equilibrium between viral RNA replication and infectious particle production (25,26). However, it is unknown whether this is a conserved co-opting mechanism in the *Orthoflavivirus* genus (which belongs also to the *Flaviviridae* family) and if so, whether these viruses modulate the composition of this complex to rewire its functions in favor of replication. Finally, considering that the 3’ NTR of vRNA is a key player of vRO morphogenesis (15), this raises the hypothesis that this viral process involves host vRNA-binding proteins prior to vRNA synthesis.

To determine whether the co-opting of these RNA-binding proteins was conserved in the *Orthoflavivirus* genus, we have assessed in this study the potential role of 10 host RBPs in ZIKV and DENV replication. We have identified insulin-like growth factor 2 mRNA-binding protein 2 (IGF2BP2) as a new host dependency factor for the ZIKV life cycle which regulates vRNA synthesis. In infected cells, IGF2BP2 associates with NS5 viral polymerase and vRNA 3’NTR. Importantly, IGF2BP2 impairs *de novo* vRO biogenesis and consistently, vRNA synthesis. Finally, ZIKV infection alters the protein and RNA interactomes of IGF2BP2. Altogether, our data support that ZIKV hijacks and remodels the IGF2BP2 ribonucleoprotein complex, to regulate vRO biogenesis and vRNA neosynthesis.

## RESULTS

### IGF2BP2 regulates ZIKV replication cycle

To identify new cellular proteins regulating ZIKV and DENV vRNA functions during viral replication, we hypothesized that these orthoflaviviruses have evolved to share conserved host co-opting mechanisms with HCV which belongs to the *Hepacivirus* genus within the *Flaviviridae* family like the *Orthoflavivirus* genus. Based on our previous work with host RBPs regulating HCV life cycle (25,26), we performed a targeted small scale RNA interference (RNAi) screening assessing the requirement of 10 host RBPs for ZIKV and DENV replication. Protein knockdown (KD) in human hepatocarcinoma Huh7.5 cells was achieved through the transduction of short-hairpin RNA (shRNA)-expressing lentiviruses, whose respective efficiencies in this cell line were previously validated (25,26). None of these shRNA had any impact on cell viability as measured by MTT assays (Fig. 1A). Two days post transduction, cells were infected with pathogenic contemporary ZIKV H/PF/2013 strain (Asian lineage), ancestral ZIKV MR766 strain (African lineage) or serotype 2 DENV2 16681s strain. Two days post-infection (dpi), infectious viral particle production was measured by plaque assays (Fig. 1B-D). The KD of DEAD-box helicases RHA (RNA helicase A, also named DHX9), DDX6, DDX21 or DDX5 had no significant impact in virus production when compared to the non-target control shRNA (shNT). In contrast, the knockdowns of IGF2BP2 and LARP1 resulted in significant decrease and increase in the titers of both ZIKV strains, respectively. The observed phenotype was specific to ZIKV since no significant impact was observed for DENV. (Fig. 1B-D).

**Figure 1:**
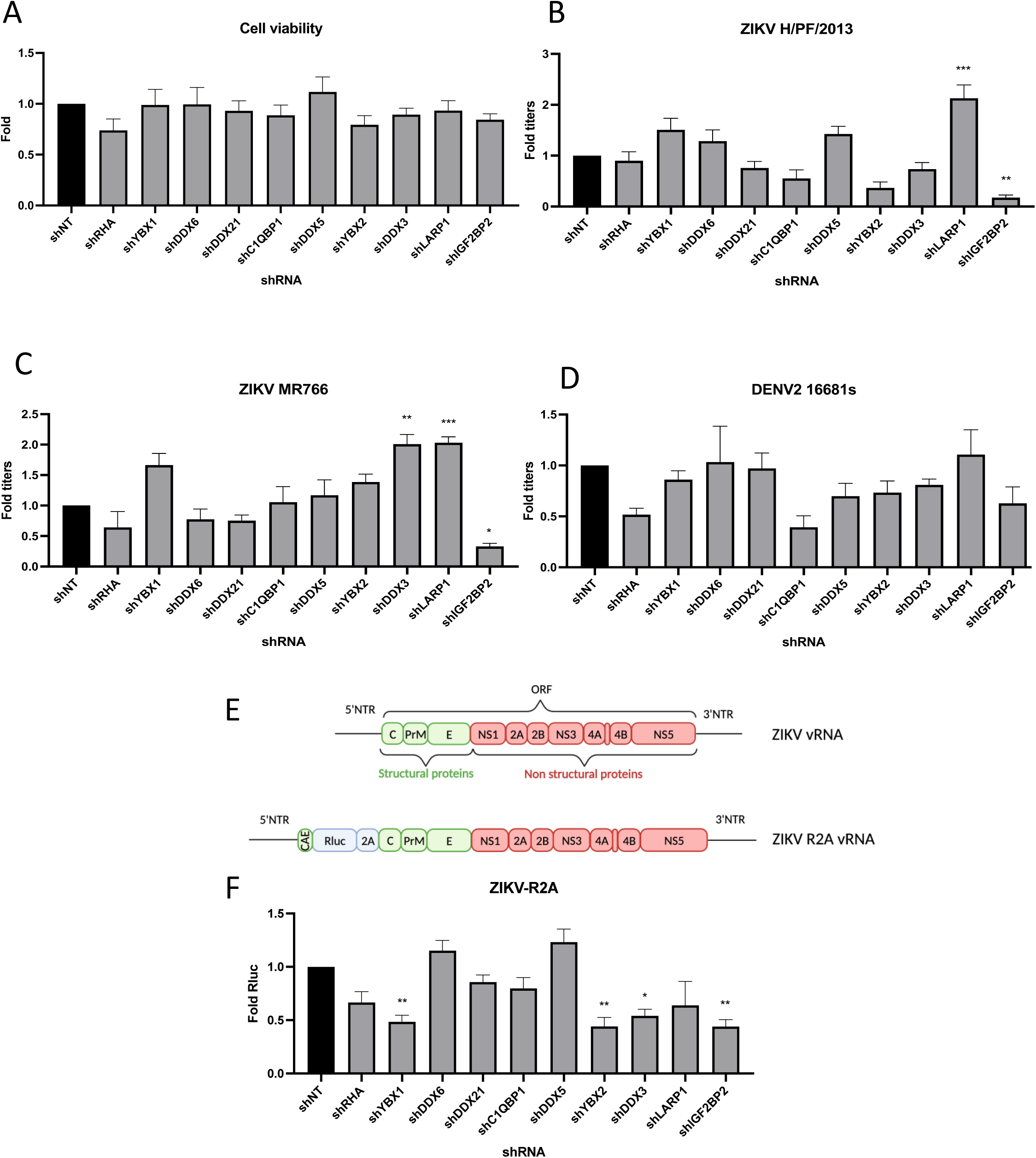
A RNAi mini-screening of RNA-binding proteins to identify host factors involved in DENV and ZIKV replications. Huh7.5 were transduced with shRNA-expressing lentiviruses at a MOI of 5-10. (A) Four days posttransduction, MTT assays were performed to evaluate cytotoxicity effect of the KD. Two days posttransduction cells were infected with either (B) ZIKV H/PF/2013, (C) ZIKV MR766, or (D) DENV2 16681s at an MOI of 0.01. 48h post infection, the production of infectious viral particles was evaluated by plaque assays. (E) Schematic of the Rluc-expressing ZIKV reporter virus (ZIKV-R2A) based on the FSS13025 isolate (Asian lineage). (F) Cells were prepared, exactly as in B-D but infected with ZIKV-R2A at a MOI of 0.001. 48h post-infection, cells were lysed and bioluminescence was measured and normalized to the control cells expressing a non-target shRNA (shNT). Means ± SEM are shown based on three to five independent experiments for each shRNA. p<0.0001; ***: p< 0.001; **: p < 0.01; * p< 0.05 (one-way ANOVA test).

To validate these phenotypes, we took advantage of ZIKV FSS13025 strain-based reporter viruses expressing Renilla luciferase (Rluc), whose activity in infected cells can be used as a read-out of overall vRNA replication (Fig. 1E) (27,28). IGF2BP2 KD resulted in a significant 60% decrease in viral replication. In contrast, LARP1 KD had no stimulatory effect on viral replication, suggesting a role of this host factor in virus assembly and/or release. Of note YBX1, YBX2 and DDX3 KD all resulted in a decrease in ZIKV replication although this did not translate into impacts in virus titers (Fig. 1B-F). We decided to focus further investigation on IGF2BP2 since it was our best candidate in terms of ZIKV replication impairment upon knockdown (Fig. 1B-C). First, western blotting and RT-qPCR with transduced Huh7.5 cell lysates validated that IGF2BP2 was efficiently knocked down upon transduction at both protein and mRNA levels (Fig. 2A-B), which correlated with a 74% decrease in ZIKV production (Fig. 2C) and reductions in the expression of viral proteins NS3, NS4A and NS5 (Fig. 2A). Comparable impairment of ZIKV titers upon IGF2BP2 knockdown was observed in other cell lines relevant for ZIKV pathogenesis, namely human immortalized astrocytes (NHA-hTERT; Fig. 2D) and cancer-derived trophoblasts (JEG-3; Fig. 2E) as well as for a third wild-type ZIKV strain in Huh7.5 cells (Fig. 3A), *i.e* ZIKV FSS13025 (Asian lineage, isolated in Cambodia in 2010).

**Figure 2:**
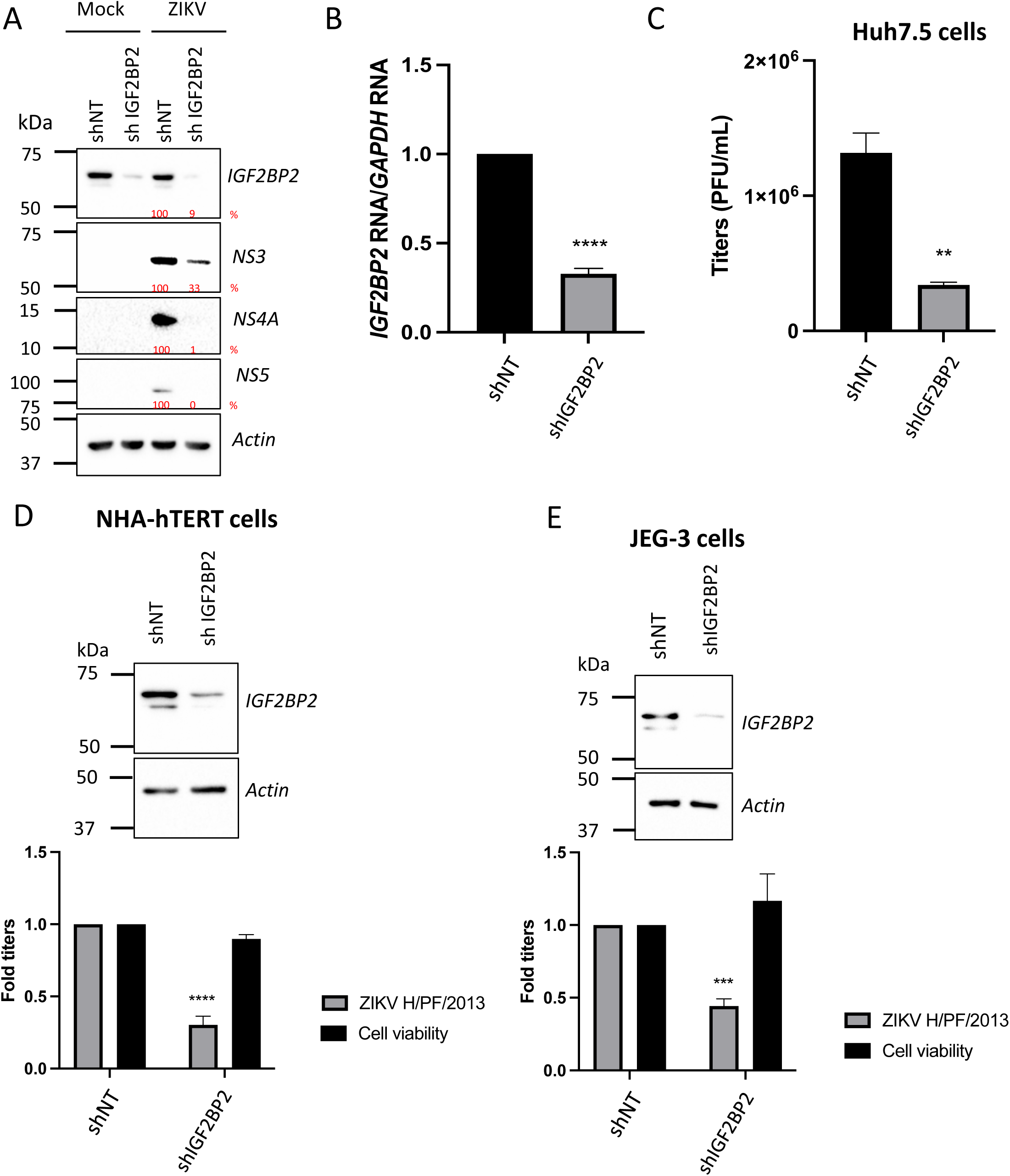
IGF2BP2 positively regulates ZIKV replication in multiple cell lines. Liver Huh7.5 (A-C), astrocytic NHA-hTERT (D) and placental JEG-3 (E) cells were transduced with shNT or shIGF2BP2 lentiviruses at a MOI of 10. Two days post transduction, cells were infected with ZIKV H/PF/2013 at a MOI between 0.01 and 1 depending on the cell line. Two days post-infection, supernatant and cells were collected. IGF2BP2 expression at the protein level (A, D, E; all cell lines) and mRNA level (B; Huh7.5 cells) were evaluated by western blotting and RT-qPCR, respectively. Cell supernatants were used for plaque assays (C-E). for NHA-hTERT and JEG-3, the supernatant and cells are collected for titration and WB respectively (E-F). MTT assays were performed to assess the cell viability in transduced NHA-hTERT and JEG-3 cells (D-E). Means ± SEM are shown based on five (D), three (C) and four (D-E) independent experiments. ****: p<0.0001; ***: p< 0.001; **: p < 0.01 (unpaired t-test).

**Figure 3:**
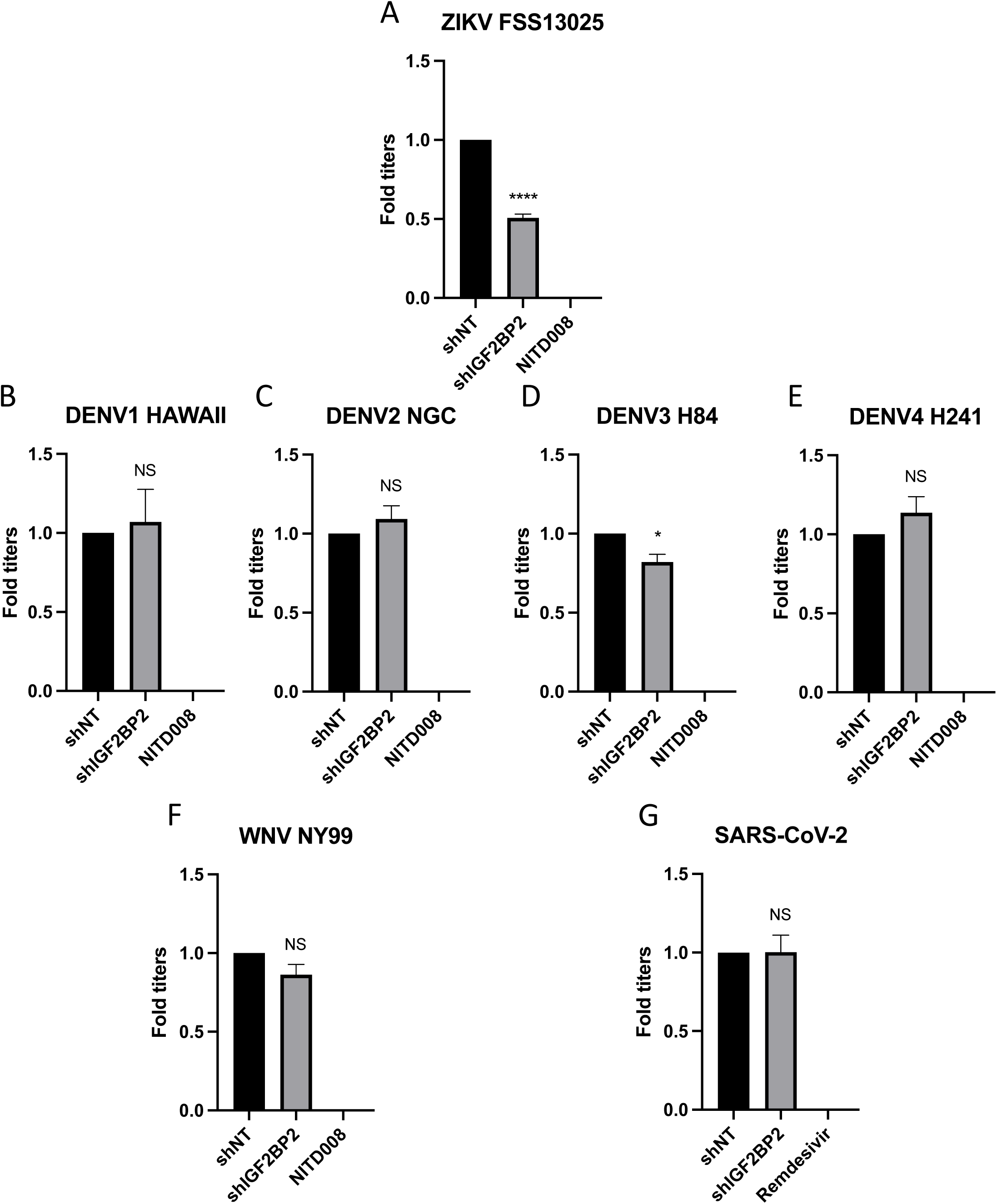
IGF2BP2 dependency is ZIKV-specific. Huh7.5 cells were transduced with shNT or shIGF2BP2 lentiviruses at a MOI of 10. Two days post-transduction cells were infected with (A) ZIKV FSS13025, (B) DENV1 HAWAII, (C) DENV2 NGC, (D) DENV3 H84, (E) DENV4 H241, (F) WNV NY99, (G) SARS-CoV-2 at a MOI of 0.1. Virus-containing cell supernatants were collected and titrated two-days post-infection by plaque assays. Treatment with RdRp inhibitors NITD008 and Remdesivir were used as positive controls of replication inhibition of orthoflaviviruses and SARS-CoV-2, respectively. Means ± SEM are shown based on three independent experiments. ****: p<0.0001; *: p < 0.05; NS: not significant (unpaired t-test).

Our initial screen showed that IGF2BP2 KD had minimal impact on DENV2 replication, suggesting a ZIKV-specific phenotype (Fig. 1D). To confirm this, we have included in our workflow four additional strains of DENV representing the four known serotypes. IGF2BP2 knockdown did not changed the production of DENV1, DENV2 or DENV4, while DENV3 replication was only slightly decreased (Fig. 3B-E). As control, treatment with NITD008, an inhibitor of flaviviral NS5 polymerase (29–32) completely abrogated viral particle production for all orthoflaviviruses tested. Finally, IGF2BP2 KD had no impact on the replication of either WNV, another orthoflavivirus (Fig.3F), or of SARS-CoV-2, another positive strand RNA virus from the *Coronaviridae* family (Fig. 3G). Overall, these data clearly show that the IGF2BP2 is a specific host dependency factor for ZIKV replication.

### IGF2BP2 redistributes to ZIKV replication complexes in infected cells

Following the observations that IGF2BP2 is a ZIKV host dependency factor, we investigated the impact of ZIKV infection on IGF2BP2 expression and subcellular distribution. We performed western blotting on extracts of either uninfected or ZIKV-infected Huh7.5 cells. IGF2BP2 expression remained unchanged at 2 and 3 days post-infection in contrast to DDX3 and DDX5 whose levels were decreased over the course of infection (Fig. 4A). Then, we investigated the localization of IGF2BP2 in infected cells using confocal microscopy. In uninfected cell, IGF2BP2 exhibited a homogenous distribution throughout the cytoplasm. In ZIKV-infected cells, an accumulation of IGF2BP2 in areas enriched for NS3 and viral double-stranded RNA (dsRNA) was observed at 2 days post-infection (Fig. 4B), correlating with a partial colocalization with NS3 and dsRNA (Manders’ coefficients of 0.45±0.04 and 0.46±0.04, respectively). dsRNA is an intermediate of vRNA replication and hence, anti-dsRNA antibodies detect replication complexes, which along with VPs, typically accumulate in a cage-like region in the perinuclear area with viral proteins required for vRNA replication step, such as NS3 and NS5 (10). This compartment is typically surrounded by NS3-positive convoluted membranes which are devoid of dsRNA (10,14,33). A similar, yet less pronounced, phenotype was observed at an earlier time point (1 dpi, Fig. S1).

**Figure 4:**
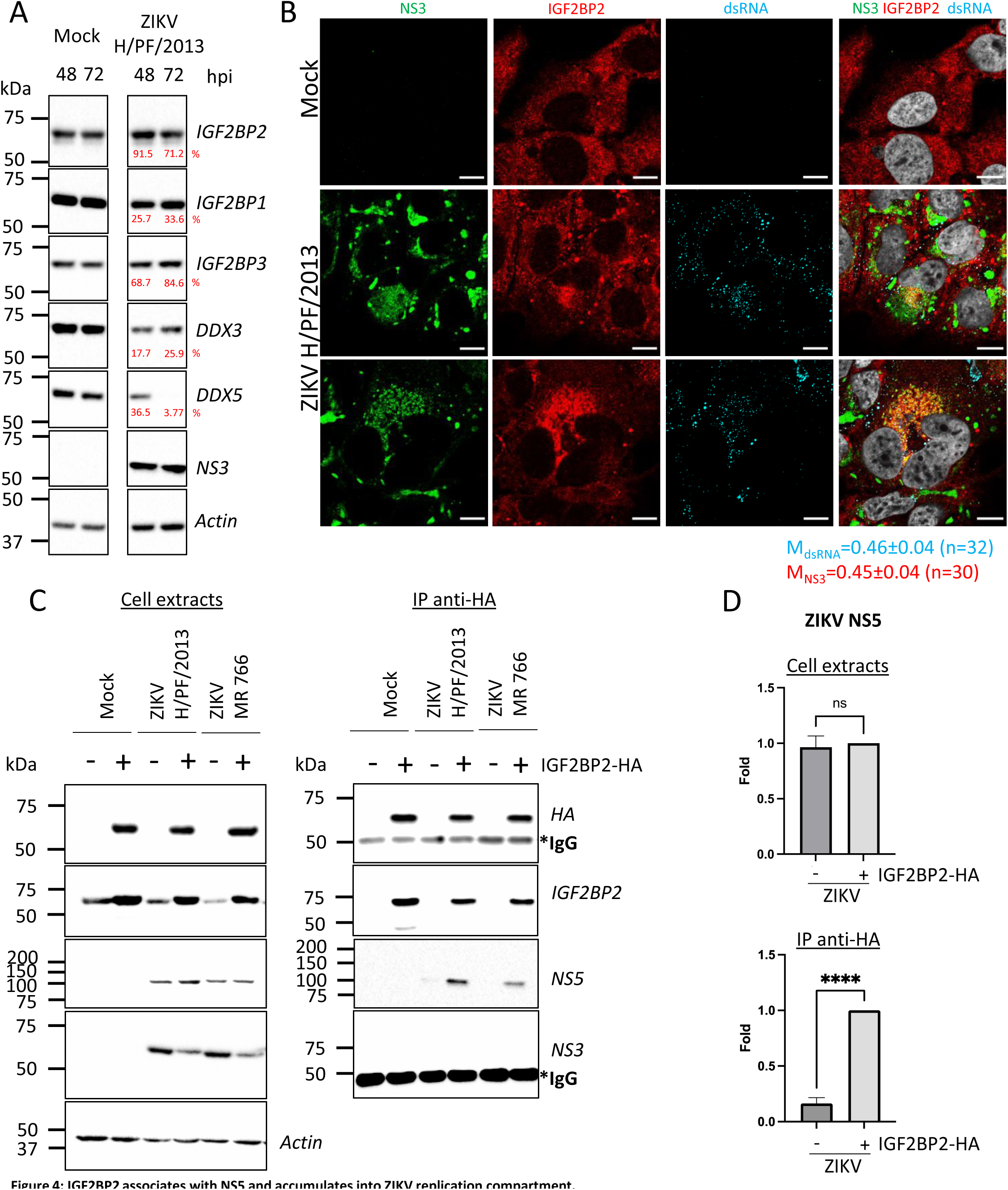
IGF2BP2 associates with NS5 and accumulates into ZIKV replication compartment. (A) Huh 7.5 cells were infected with ZIKV H/PF/2013 at a MOI of 5. Cells were collected at 48-and 72-hours postinfection (hpi). Cell extracts were prepared and analyzed by western blotting using the indicated antibodies. Actin-normalized protein signals are shown. (B) Huh7.5 cells were infected with ZIKV H/PF/2013 with a MOI of 10 or left uninfected. Two days post-infection, cells were fixed, immunolabelled for the indicated factors, and imaged by confocal microscopy. Scale bar=10 m). The Manders’ coefficient (mean ± SEM) representing the fraction of dsRNA (cyan) and NS3 (red) signals overlapping with IGF2BP2 signal is shown (n=number of cells). (C) Co-immunoprecipitation assays using HA antibodies were performed with extracts from Huh7.5 cells stably expressing IGF2BP2-HA (+) or control transduced cells (-) which were infected with ZIKV at a MOI of 10 for 2 days. Purified complexed were analyzed for their protein content by western blotting. (D) Means of quantified NS5 signals from C (normalized to actin (extracts) or IGF2BP2 (IP)) ± SEM are shown based on nine independent experiments. ****: p<0.0001; ns: not significant (unpaired t-test).

The relocalization of IGF2BP2 into the replication compartment raised the hypothesis that this protein interacts with viral proteins involved in vRNA replication. To test this, we generated Huh7.5 cells stably expressing HA-tagged IGF2BP2 to subsequently perform co-immunoprecipitation assays (Fig. 4C). As control, we first confirmed that this tagged recombinant protein, similarly to endogenous IGF2BP2, also redistributed to the dsRNA-positive area in ZIKV infected cells (Fig. S2). We next performed anti-HA immunoprecipitations with extracts from uninfected and ZIKV infected cells. Western blot analysis of immunopurified complexes showed that IGF2BP2-HA specifically associates with ZIKV NS5 polymerase but not with NS3 protease/helicase (Fig. 4C-D). Overall, these data show that ZIKV infection induces the physical recruitment of IGF2BP2 to the replication compartment, and further suggest that IGF2BP2 might contribute to vRNA synthesis through interactions with NS5 polymerase.

### IGF2BP2 associates with ZIKV RNA

Considering that IGF2BP2 associates with NS5 in the replication compartment and that it possesses RNA-binding activities, we evaluated whether IGF2BP2 interacts with ZIKV vRNA. IGF2BP2, along with IGF2BP1 and IGF2BP3 paralogues are highly conserved oncofetal RNA-binding proteins (RBPs) belonging to the Insulin-like growth factor 2 mRNA-binding protein family, which typically regulate the post-transcriptional regulation of cellular RNAs. With their roles in the splicing, transport, translation, and stabilization of a wide variety of RNAs (34,35), IGF2BP1, 2 and 3 are involved in numerous cellular function, such as differentiation, migration, metabolism and proliferation (34,35). IGF2BP2 is composed of six canonical RNA-binding domains, namely two N-terminal RNA-recognition motifs (RRM) and four C-terminal human heterogeneous nuclear ribonucleoprotein (hnRNP)-K homology (KH) domains (36–38).

To assess a potential association between IGF2BP2 and ZIKV vRNA, we first performed protein immunostaining coupled to single RNA molecule fluorescence *in situ* hybridization (FISH) using signal-amplified DNA branched probes to detect IGF2BP2 and ZIKV RNA, respectively. ZIKV RNA was specifically visualized since no FISH signal was detected in uninfected cells while in ZIKV-infected cells, vRNA exhibited a punctate distribution (Fig. 5A, Fig. S3). Co-staining of vRNA and IGF2BP2 revealed their partial colocalization (Fig. 5A, white arrows, mean Manders’ coefficient=0.28±0.02 (n=31)), which strongly suggests that IGF2BP2 associates with the ZIKV RNA genome.

**Figure 5:**
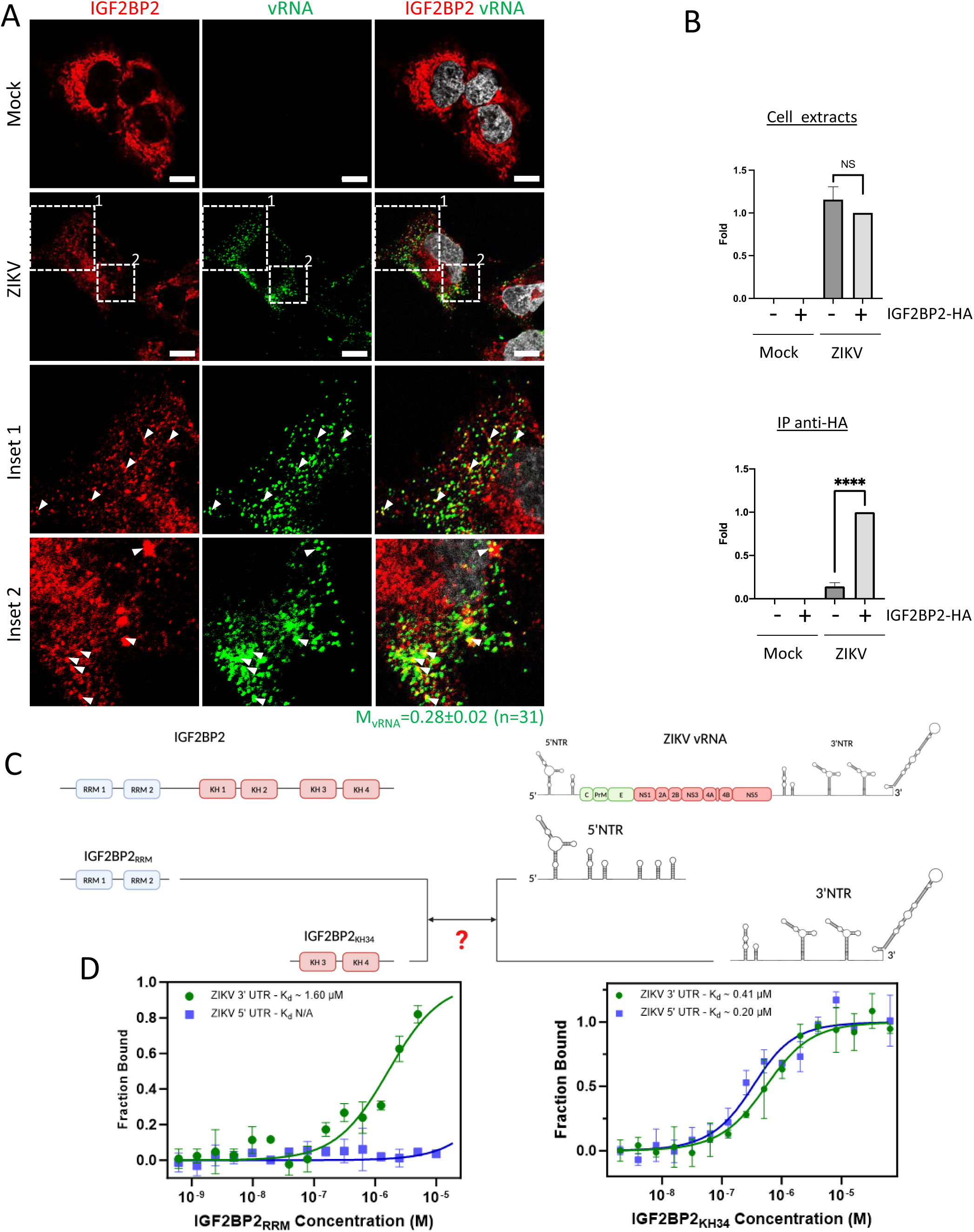
IGF2BP2 interacts with ZIKV vRNA. (A) FISH and IGF2BP2 immunostaining were performed using Huh 7.5 cells which were infected for 2 days with ZIKV (MOI = 10) or left uninfected, The Manders’ coefficient (mean ± SEM) representing the fraction of vRNA signal overlapping with IGF2BP2 signal is shown (n=number of cells). (B) Huh7.5 cells expressing IGF2BP2-HA and control cells were infected with ZIKV H/PF/2013 at a MOI of 10, or left uninfected. Two days later, cell extracts were prepared and subjected to anti-HA immunoprecipitations. Extracted vRNA levels were measured by RT-qPCR. Means ± SEM are shown based on three independent experiments. ****: p<0.0001; NS: not significant (unpaired t-test). (C) IGF2BP2 recombinant proteins containing either the two RRM or KH3 and KH4 domains were produced in bacteria and purified. In parallel ZIKV 5’ NTR and 3’ NTR were synthesized by in vitro transcription. (D) Combination of truncated IGF2BP2 proteins and either ZIKV 5’ NTR (blue squares) or ZIKV 3’ NTR (green circles) were used for in vitro binding assays using microscale thermophoresis.

To test this, we performed anti-HA co-immunoprecipitation assays using extracts of Huh7.5 cells as in Fig. 4C. Immune complexes were subjected to RNA purification and subsequently to RT-qPCR to detect viral genome (Fig. 5B). While the overexpression of IGF2BP2 had no impact on total vRNA levels, we detected a highly significant enrichment of vRNA in purified IGF2BP2-HA complexes compared to the negative specificity control (no IGF2BP2-HA expression). This demonstrates that vRNA and IGF2BP2 are part of the same RNP complex. To further investigate if IGF2BP2 can interact with ZIKV vRNA in a direct manner, we have performed *in vitro* binding assays using microscale thermophoresis (MST) with different regions of ZIKV vRNA and IGF2BP2 protein. We have focused our analysis on vRNA 5’ NTR and 3’ NTR, which are highly structured and are absolutely required for vRNA synthesis and translation, notably through interactions with viral and host factors (for a review see (8)). Moreover, it has been shown that IGF2BP2 preferentially binds to 3′ and 5′ untranslated regions of mRNAs (39). ZIKV RNA 3’ NTR and 5’ NTR were synthetized by *in vitro* transcription. In parallel recombinant IGF2BP2 N-terminal and C-terminal moieties containing either the two RRM (IGF2BP2_RRM_) or both KH3 and KH4 domains (IGF2BP2_KH34_), respectively were produced in bacteria and subsequently purified (Fig. 5C). MST revealed that IGF2BP2_KH34_ binds both ZIKV 5’ NTR and 3’ NTR with high affinity with respective K_d_ of 203 nM ± 51 and 418 nM ± 49, respectively (Fig. 5D). In stark contrast, IGF2BP2_RRM_ specifically associated with ZIKV 3’ NTR with a lower affinity (K_d_ = 1,598 nM ± 257) but this interaction was highly specific since no binding was detected between this recombinant protein and ZIKV 5’ NTR. It is noteworthy to mention that we could not assess higher IGF2BP2_RRM_ concentrations than 10 μM to reach binding saturation because of protein aggregation. Overall, these data demonstrate that IGF2BP2 directly and specifically interacts with vRNA in infected cells.

### ZIKV regulates vRNA replication

The fact that IGF2BP2 associates with both ZIKV NS5 RdRp and vRNA led us to hypothesize that IGF2BP2 is involved in the vRNA replication step of ZIKV life cycle. To test this, we took advantage of reporter ZIKV subgenomic replicons based on the H/PF/2013 genome (ZIKV sgR2A; Fig. 6A)(40). This engineered genome has been deleted for the coding sequence of structural proteins and expresses Rluc in frame with the NS1-NS5 polyprotein. When *in vitro*-transcribed genomes are introduced in cells by electroporation, they autonomously replicate but neither virus assembly nor entry occur because of the lack of structural proteins. Hence, the luciferase activity in transduced Huh7.5 cells was used as a read-out of viral RNA replication two days post-electroporation. Strikingly, ZIKV sgR2A replication was attenuated in IGF2BP2 KD, with a significant 40% decrease (Fig. 6B). To rule out that this phenotype was due to a potential defect in vRNA translation, we used mutated Rluc-expressing ZIKV sub-genomes which express a defective NS5 RdRp and do no replicate (ZIKV sgR2A GAA; Fig. 6A, B). Hence, the Rluc activity detected at 4 hours post-electroporation entirely relies on viral RNA translation. IGF2BP2 knockdown did not have any significant impact on Rluc activity in Huh7.5 cells transfected with either sgR2A or sgR2A GAA (Fig. 6C). These data show that IGF2BP2 positively regulates ZIKV genome replication (but not its translation), which is most likely mediated through its interactions with NS5 and vRNA.

**Figure 6:**
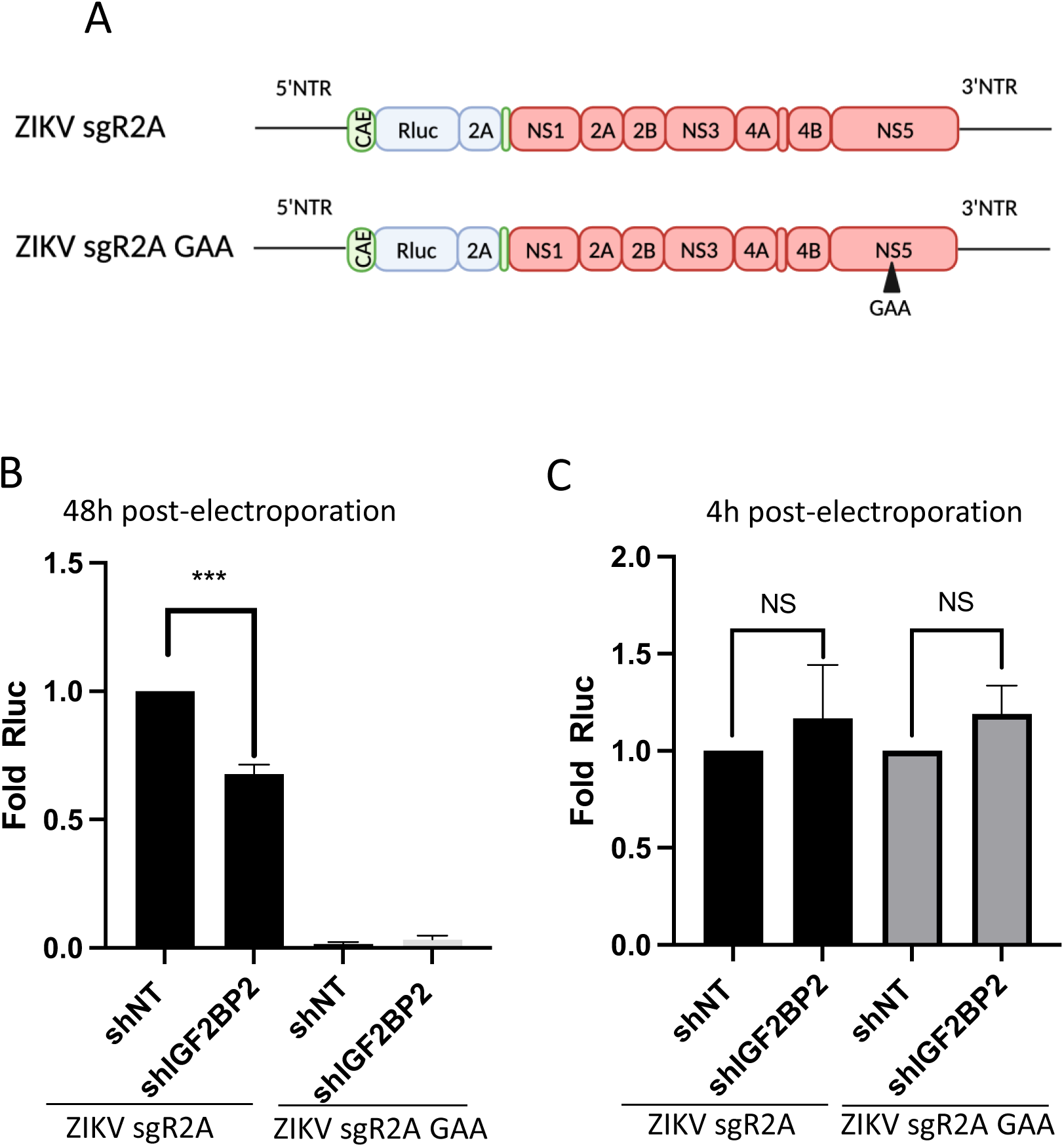
IGF2BP2 regulates the replication of ZIKV vRNA. (A) Schematic representation of reporter ZIKV H/PF/2013 sub-genomic replicons (sgR2A) and replication-deficient genomes because of mutations in NS5 RdRp sequence (sgR2A GAA). (B-C) Huh7.5 were transduced with shRNA-expressing lentiviruses and subjected to electroporation with in vitro transcribed sgR2A or sgR2A GAA RNAs two days later. In-cell bioluminescence was measured (B) 48 or (C) 4 hours post-electroporation and normalized to the shNT control condition. In C, the luciferase activity was normalized to the transfection efficiency i.e., the Rluc activity at 4h post-electroporation. Means ± SEM are shown based on four independent experiments. ***: p<0.001; NS: not significant (unpaired t-test).

### ZIKV infection modulates the interactions between IGF2BP2 and its endogenous mRNA ligands

Since IGF2BP2 associates with NS5 and vRNA and accumulates in the viral replication compartment, we hypothesized that ZIKV infection induces a remodeling of IGF2BP2 RNP notably regarding its content in endogenous mRNA partners. To test this, we performed immunopurifications of IGF2BP2 complexes followed by RT-qPCR as in Figs. 4C and 5B in order to determine the relative abundance of three known IGF2BP2 mRNA ligands (namely *TNRC6A*, *PUM2* and *CIRBP*)(41) in both control and ZIKV-infected cells. *TRNC6A, PUM2* and *CIRBP* mRNAs were selected because ZIKV is known to alter their N6-adenosine methylation status (42), an epitranscriptomic RNA modification that increases the affinity for IGF2BP2, a known “m6A reader”(41). As expected, we could specifically detect these mRNA in purified IGF2BP2 complexes in uninfected cells (Mock+IGF2BP2-HA condition; Figs. 7A-C). Interestingly, the levels of co-immunoprecipitated *TNRC6A* and *PUM2* mRNAs cell were significantly decreased when Huh7.5 were infected with ZIKV while total mRNA levels remained unchanged (Fig. 7A-B), suggesting that ZIKV induced a loss of binding between IGF2BP2 and these endogenous mRNAs. In contrast, no significant changes in IGF2BP2 association were observed for *CIRBP* mRNA in either condition although it is noteworthy that total levels were increased in total extracts when IGF2BP2 was overexpressed (Fig.7C). These results show that ZIKV infection modifies the interaction between IGF2BP2 and specific endogenous mRNAs and is consistent with the notion of a virus-induced remodeling of IGF2BP2 RNP.

**Figure 7:**
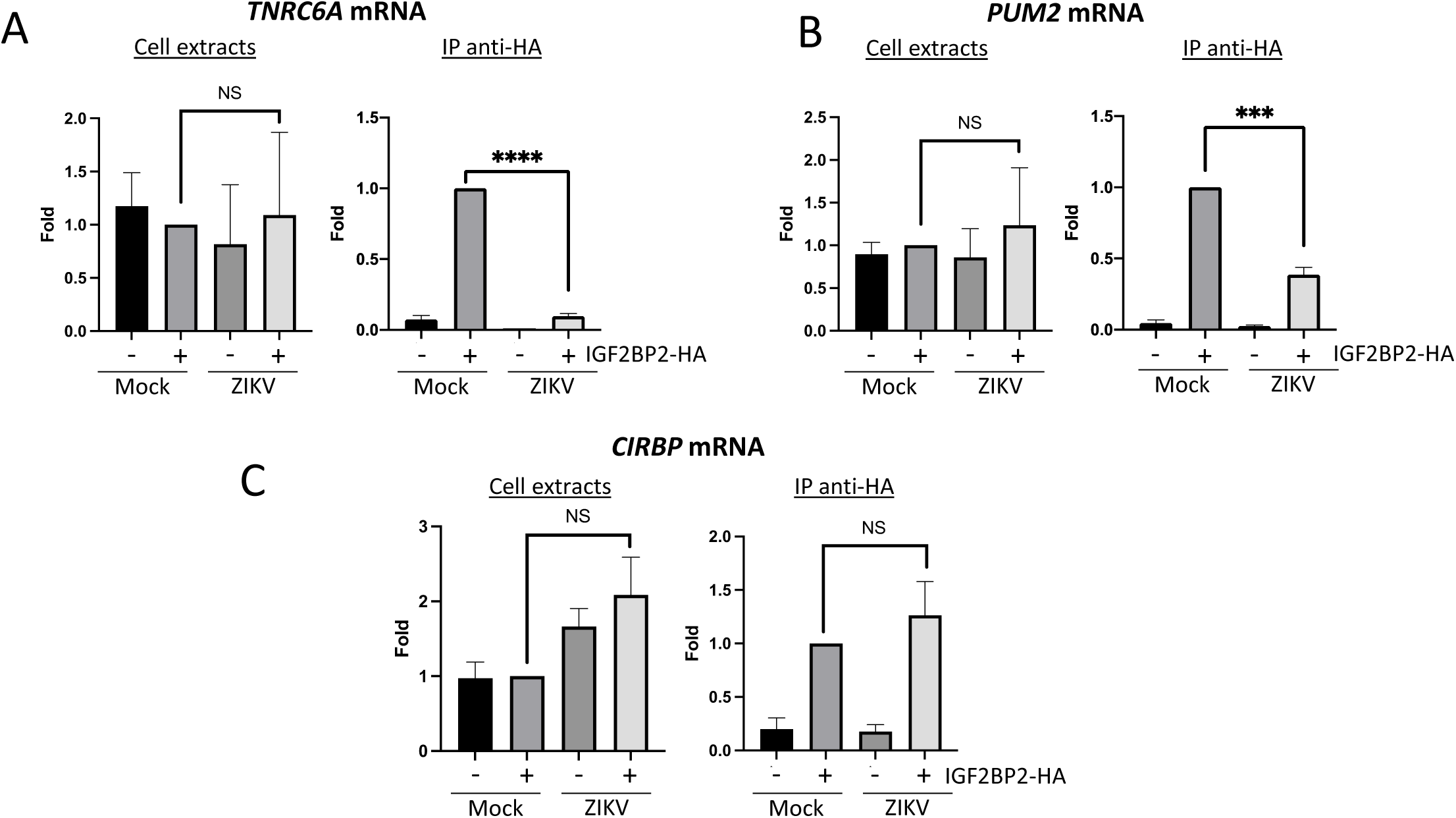
ZIKV infection decreases the interaction between IGF2BP2 and several of its mRNA endogenous ligands. Huh7.5 cells stably expressing IGF2BP2-HA (+) and control cells (-) were infected with ZIKV H/PF/2013 at a MOI of 10, or left uninfected. Two days later, cell extracts were prepared and subjected to anti-HA immunoprecipitations. Extracted (A) TNRC6A, (B) PUM2, and (C) CIRBP mRNA levels were measured by RT-qPCR. Means ± SEM are shown based on three independent experiments. ****: p<0.0001; ***: p<0.001; NS: not significant (unpaired t-test).

### ZIKV alters IGF2BP2 proteo-interactome in infected cells

Next, to assess our hypothesis of a ZIKV-remodeled IGF2BP2 RNP, we investigated whether ZIKV infection changes the protein interaction profile of IGF2BP2. We first performed western blotting on purified IGF2BP2-HA samples (prepared as in Figs. 4C and 5B) to detect IGF2BP1, IGF2BP3 and YBX1 which are known IGF2BP2 partners (26,37,43). Comparable amounts of either partner were specifically co-purified with IGF2BP2-HA (Figs. 8A-D, S6A and S7C) indicating that ZIKV infection does not significantly change their association. As expected from results shown in 4C, the viral RdRp NS5 was readily detected in this complex. Interestingly, confocal microscopy showed ZIKV infection induced an accumulation of IGF2BP1, IGF2BP3 and YBX1 in the replication compartment as for IGF2BP2 with partial colocalization with dsRNA and NS3 (Figs. S4, S5C-D), which supports that ZIKV physically co-opts an RNP containing these fours RBPs. This phenotype was specific to these host factors as it was not observed for the RNA-binding proteins DDX5 and LARP1 (Fig. S5A-D). Consistent with the fact that these proteins (including NS5) belong to a ribonucleoprotein complex containing IGF2BP2, their interaction with IGF2BP2 was RNA-dependent since they were barely detectable in the immunoprecipitates when the cell extracts were treated with RNase A prior to anti-HA pull-down (Fig. S6) To globally evaluate changes in the IGF2BP2 interactome upon infection, we analyzed the protein composition of IGF2BP2-HA complexes immunopurified from uninfected cells or infected with ZIKV or DENV by mass spectrometry. We identified 527 proteins which specifically interacted with IGF2BP2 (Fig. S7A-B; Table S1). The abundance of over 86% of the proteins in IGF2BP2-HA complexes (455 proteins), including IGF2BP1, IGF2BP3 and YBX1 remained unchanged upon either infection (as expected from Fig. 8A-D). In contrast, in ZIKV-infected cells, the interactions of IGF2BP2 with 40 and 22 proteins were either decreased or increased, respectively, with 52 of these changes being specifically observed only in ZIKV-infected cells (Fig. 9A; Fig. S7A). Gene ontology analysis of the 62 partners whose interaction with IGF2BP2 was altered during ZIKV infection, revealed a high enrichment of biological processes related to mRNA splicing (Fig. 9B). The interactome tree of these 62 IGF2BP2 protein partners generated with STRING database highlighted two major clusters regulated by ZIKV (Fig. 9C). In line with our gene ontology analysis, one of these clusters comprised proteins involved in mRNA splicing for most of which interaction with IGF2BP2 was significantly decreased during ZIKV infection (red circles). The second identified cluster was related to ribosome biogenesis.

**Figure 8:**
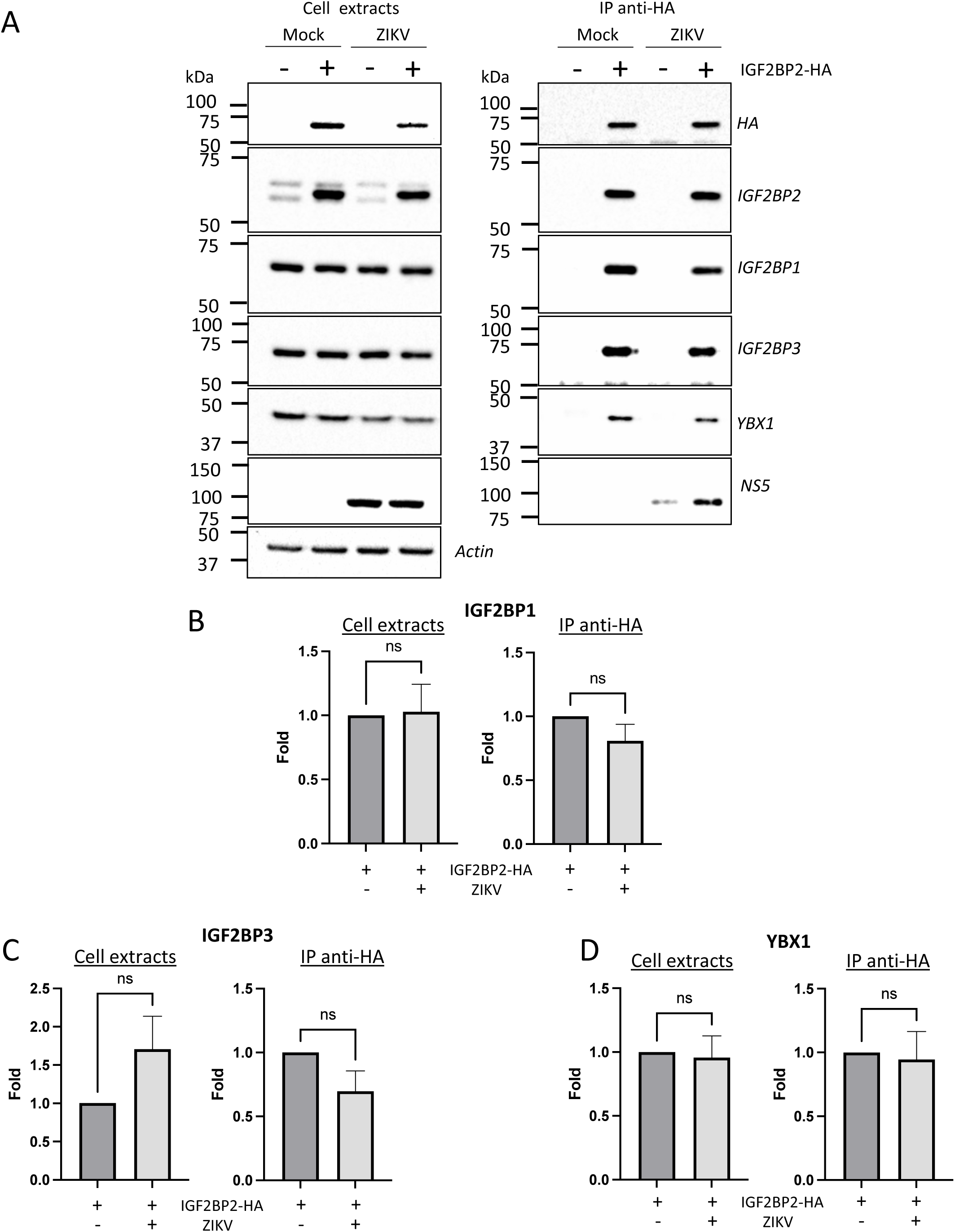
ZIKV infection does not significantly impact the association of IGF2BP2 with IGF2BP1, IGF2BP3 and YBX1. Huh7.5 cells stably expressing IGF2BP2-HA (+) and control cells (-) were infected with ZIKV H/PF/2013 at a MOI of 10, or left uninfected. Two days later, cell extracts were prepared and subjected to anti-HA immunoprecipitations. (A) Purified complexes were analyzed by western blotting for their content in the indicated proteins. IGF2BP1 (B), IGF2BP3 (C) and YBX1 (D) levels were quantified and means of protein signals (normalized to actin (extracts) and IGF2BP2 (IP)) ± SEM are shown based on six to eight independent experiments. ns: not significant (unpaired t-test).

**Figure 9:**
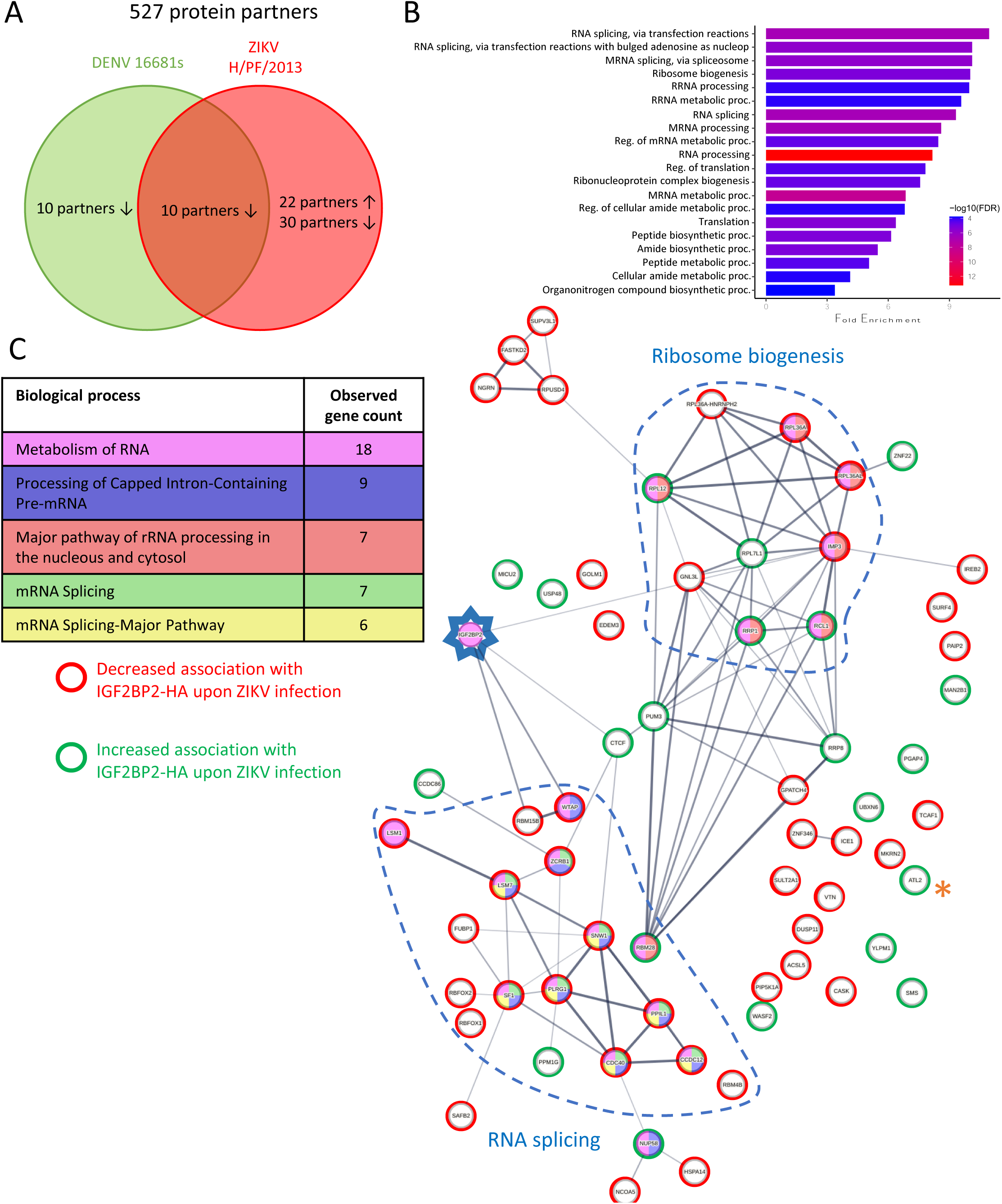
ZIKV infection alters the IGF2BP2 proteo-interactome. Huh7.5 cells expressing IGF2BP2-HA and control cells were infected with ZIKV H/PF/2013, DENV2 16681s, or left uninfected. Two days later, cell extracts were prepared and subjected to anti-HA immunoprecipitations. Resulting complexes were analyzed by quantitative mass spectrometry. (A) Venn diagram depicting the overlap between IGF2BP2 partners modulated by ZIKV and/or DENV infections. (B) Gene Ontology (GO) biological process analyses of the IGF2BP2 interactions which were impacted upon ZIKV infection. (C) Interaction tree of the 62 IGF2BP2 interactions modulated by ZIKV infection (generated with STRING online resource). The red and green circles identify the partners of the STRING network whose association with IGF2BP2 is decreased and increased during infection, respectively. The biological process analysis generated by STRING is also shown.

One of the best partners specifically modulated by ZIKV and/or DENV was Atlastin 2 (ATL2) whose interaction with IGF2BP2 was the most significantly increased upon ZIKV infection (p-value = 10^-5.7^; Table S1; highlighted with * in Fig. 9C and S4B). ATL2 is an ER-shaping protein (44) which was reported to be involved in the formation of orthoflavivirus vesicle packets (18). We validated this ZIKV-specific phenotype by co-immunoprecipitation assays (Figs. S6A, S7C-D) in which we could detect a 14-fold increase in IGF2BP2-HA/ATL2 interaction when cells were infected with ZIKV compared to uninfected or DENV-infected samples. As an additional specificity control, we did not detect any interaction between ATL2 and HA-tagged VCP (Fig. S7C), an ER quality control protein which was shown to be physically recruited into DENV and ZIKV replication compartments (17,33), ruling out that ATL2/IGF2BP2 interaction is simply due to the presence of ATL2 within remodeled ER or because of non-specific binding to the HA-tag. Altogether, these data show that ZIKV infection specifically alter the composition of IGF2BP2 RNP complex.

### IGF2BP2 is involved in the biogenesis of ZIKV replication organelles

The fact that ZIKV infection promotes IGF2BP2 association with ATL2 led us to hypothesize that IGF2BP2 regulates vRNA replication by contributing to ZIKV replication organelle biogenesis. Since IGF2BP2 KD decreases vRNA replication and thus, indirectly reduces the overall synthesis of viral proteins that drive vRO formation, evaluating this hypothesis in infected cells was not possible. To tackle this challenge, we took advantage of a recently described plasmid-based system (named pIRO-Z)(15,16) which induces vROs *de novo* in transfected cells in a replication-independent manner (Fig. 10A). This plasmid allows the cytoplasmic transcription of NS1-NS5 polyprotein-encoding mRNA under the control of T7 RNA polymerase which is stably overexpressed in Huh7-derived Lunet-T7 cells.

**Figure 10:**
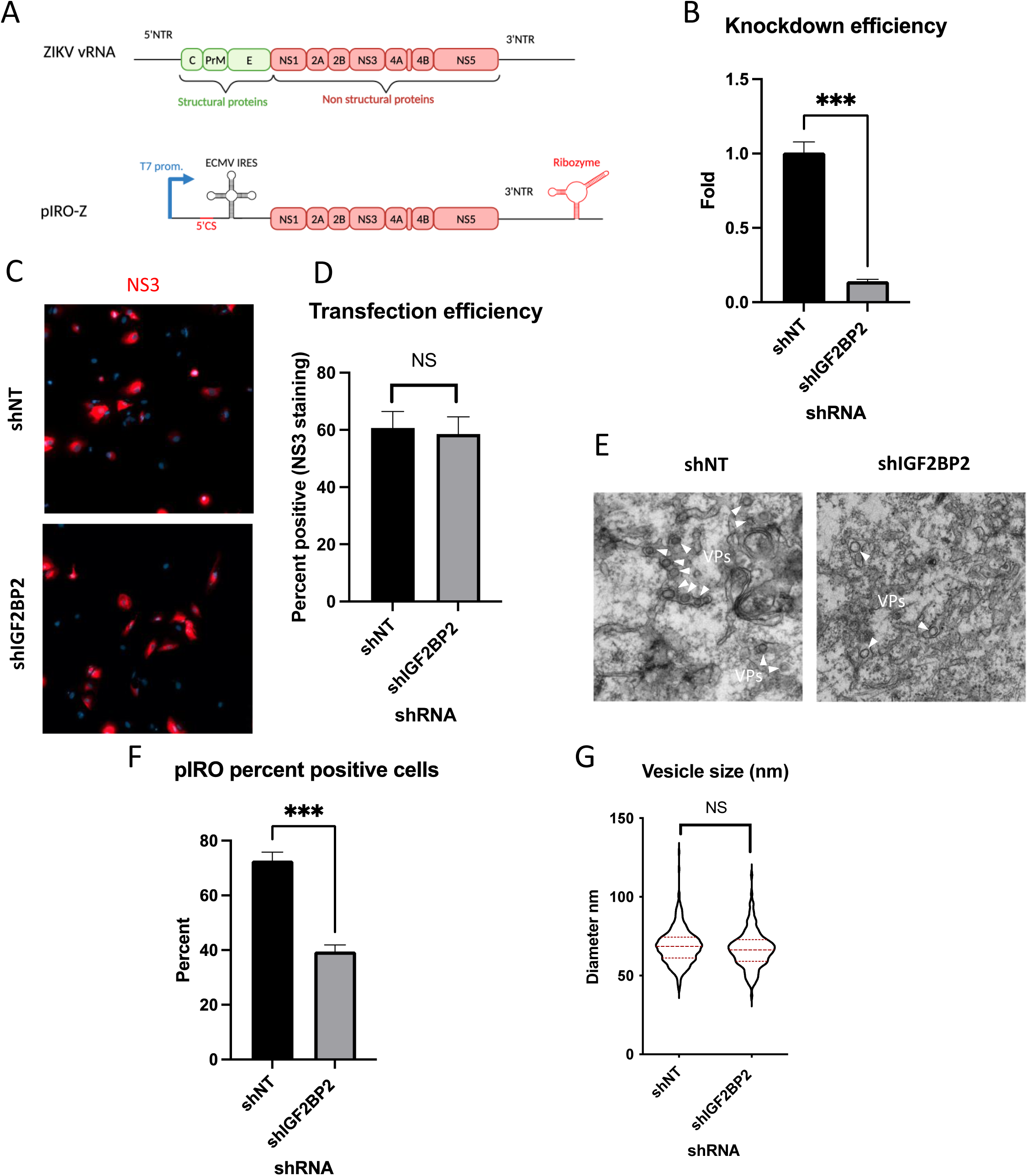
IGF2BP2 regulates the biogenesis of ZIKV replication organelles. (A) Schematic representation of the pIRO system. Upon transfection in cells expressing the T7 RNA polymerase, this plasmid allows the cytoplasmic transcription of NS1-NS5 polyprotein under the control of T7 promoter, in a ZIKV replication-independent manner. NS1-5 polyprotein synthesis is under the control of ECMV IRES. The presence of both ZIKV 3’ NTR and 5’ cyclization sequence (5’ CS) is required for efficient VP induction. Finally, the activity of HDV ribozyme ensures that the 3’ terminus of the RNA is similar to that of vRNA genome. Huh7-Lunet-T7 were transduced with shRNA-expressing lentiviruses at a MOI of 5-10. Two days later, transduced cells were transfected with pIRO-Z plasmid. Sixteen hours later, cells were analyzed for (B) IGF2BP2 mRNA levels by RT-qPCR to measure knockdown efficiency, (C-D) transfection efficiency by confocal imaging of NS3-labeled cells, and (E) for VP content by transmission electron microscopy. Electron micrographs were used to measure (F) the percentage of cells with VPs and (G) the diameter of VPs in each condition. ***: p<0.001; NS: not significant (unpaired t-test).

Lunet-T7 cells were transduced with shNT/shIGF2BP2-expressing lentiviruses and subsequently transfected with pIRO-Z. The efficiency of IGF2BP2 knockdown was controlled by RT-qPCR (Fig. 10B). Confocal microscopy of NS3-immunolabeled cells confirmed that the transfection efficiencies were comparable between all conditions (Fig. 10C-D). When imaged by electron microscopy, VPs were detected in 70% of the shNT-transduced cells. In contrast, IGF2BP2 knockdown resulted in a decrease in the proportion of cells containing vROs (Fig. 10E-F) while the diameter of VPs remained unchanged (Fig. 10G). These results demonstrate that IGF2BP2 plays a role in the biogenesis of ZIKV VPs independently from vRNA synthesis.

## DISCUSSION

In this study, to identify new host factors that regulate viral processes involving the orthoflavivirus vRNA, we have assessed 10 RNA-binding-proteins which were previously reported to regulate the life cycle of HCV, an hepacivirus belonging to the same *Flaviviridae* family as DENV and ZIKV (25,26). For instance, YBX1, IGF2BP2, LARP1, DDX6, C1QBP1 were reported to positively regulate HCV RNA synthesis while inhibiting the production of infectious viral particles. Our RNAi mini-screen assessing the impact of gene depletion on DENV or ZIKV infection revealed that overall, such host factor-mediated regulations are not fully conserved across the *Flaviviridae* family. Only LARP1 knockdown reduced ZIKV replication while it increased virus production as observed for HCV (26). A comparable trend was observed for YBX1 although the differences were not significant. In case of DENV, none of the tested RNA-binding proteins increased infectious particle production. The fact that the phenotypes did not fully mirror those we have reported for HCV might reflect the differences in the architecture of the vROs induced by orthoflaviviruses (*i.e.*, invaginated vesicles in the ER (10,11,45–47)) compared to that of HCV double-membrane vesicles which result from ER protrusions (48). The best observed phenotype in our RNAi mini-screen was observed with IGF2BP2 knockdown which inhibited the replication of ZIKV but not that of other tested orthoflaviviruses or SARS-CoV-2, another positive-strand RNA virus from the *Coronaviridae* family. We further observed that ZIKV induced a relocalization of IGF2BP2 to the replication compartment in which it associates with NS5 and vRNA. Consistently, IGF2BP2 knockdown decreased the efficiency of vRNA synthesis (but not its translation) correlating with an impairment in VP formation (Figs. 6 and 10). This highlights that IGF2BP2 is an important regulator of the vRNA replication step of ZIKV life cycle. Of note, the overexpression did not result in an increase of vRNA abundance. We believe that this is because endogenous IGF2BP2 is highly expressed in cancer cells such as the Huh7.5 cells used here and is presumably not limiting for viral replication in this system (49–53).

Early in the ZIKV replication cycle, newly produced viral nonstructural proteins induce ER alterations to form VPs which will host the vRNA synthesis machinery including NS5 RNA polymerase (54). However, the molecular mechanisms governing VP morphogenesis remain poorly understood, mostly because it is experimentally challenging to discriminate phenotypes related to vRO morphogenesis from those resulting from perturbations in vRNA replication which reduce viral protein input and hence, VP abundance indirectly. In this study, using a recently engineered replication-independent VP induction system (15,16), we identified IGF2BP2 as a novel regulator of ZIKV *de novo* VP biogenesis. Interestingly, IGF2BP2 associates with ZIKV RNA 3’ NTR and ATL2, which are both co-factors of VP formation (15,18), suggesting that they all regulate this process as part of the same RNP complex. *In vitro* RNA binding assays showed that IGF2BP2 can interact directly and specifically with vRNA 3’NTR via its RRM domains. A recombinant protein comprising the third and fourth KH domains of IGF2BP2 also bound 3’ NTR. However, this interaction appeared less specific since a comparable affinity was observed for the 5’ NTR *in vitro*. Besides, we cannot exclude that IGF2BP2 binds additional vRNA regions in the coding sequence, which was not tested in this study. Interestingly, it has been demonstrated that the KH domains of IGF2BPs mediate the binding to *N^6^*-methylated adenosines in mRNA (41), a modification that has been detected within the RNA genome of ZIKV and other *Flavividae* viruses, including in the 3’ NTR (55–58). Since IGF2BP2 is a “m6A reader” through KH3 and KH4 (41,59), it is highly plausible that this RNA modification contributes to the vRNA binding specificity and the proviral roles of this host factor. Considering that the binding of KH3 and KH4 of IGF2BP1 paralogue to RNA can induce conformational changes of the RNA (60), it will be interesting to evaluate whether IGF2BP2 co-opting by replication complexes (most likely via NS5) contributes to the structural switches and/or secondary and tertiary structures of vRNA controlling its fate during the life cycle.

To study the impact of ZIKV infection on IGF2BP2 association with endogenous RNAs, we decided to focus our analysis on three known IGF2BP2 mRNA ligands namely *CIRBP*, *TNRC6A* and *PUM2*. We chose to focus on those mRNA because: i) ZIKV, in addition to other *Flaviviridae* viruses (DENV, WNV and HCV) alter their N^6^-adenosine methylation status during infection (55–57), and ii) IGF2BP2 is a well described “m^6^A reader” which regulates the fate of bound RNAs (41,59) . More specifically, it has been shown that ZIKV infection decreases m^6^A content of *CIRBP* mRNA and increases the one of *PUM2* and *TNRC6A* mRNAs. Unexpectedly, we observed a decreased interaction between these overmethylated mRNAs and IGF2BP2 while IGF2BP2/*CIRBP* mRNA association remained unchanged (Fig. 7). This counterintuitive observation might be due to additional layers of IGF2BP2 regulation such as potential virus-dependent posttranslational modifications. Indeed, the mTOR complex 1 (mTORC1) was reported to phosphorylate two residues of IGF2BP2 (Ser162/Ser164) which are located between the second RRM domain and the first KH domain (61). The simultaneous phosphorylation of these two serine residues strongly enhances the interaction between IGF2BP2 and IGF-II leader 3 mRNA 5’UTR. Interestingly, our preliminary mass spectrometry analysis of IGF2BP2 phosphopeptides abundance indicate that ZIKV decreases phosphorylation of Ser162, but not that of Ser164. Moreover, several studies reported that ZIKV infection inhibits mTORC1 signaling (62,63). Thus, it is tempting to speculate that ZIKV, through the mTOR pathway, regulates the affinity of IGF2BP2 for endogenous mRNAs by modulating its phosphorylation status. Our interactome analysis revealed global changes in the composition of IGF2BP2 RNP with 62 cellular proteins whose association with IGF2BP2 was altered during ZIKV infection. Most notably, ZIKV infection decreased IGF2BP2 interaction with 15 proteins of the mRNA splicing machinery. Interestingly, there is evidence that IGF2BP2 regulates mRNA splicing (39) in addition to its most characterized functions in mRNA stability and translation. Furthermore, several studies reported that ZIKV infection alters mRNA splicing (64–66). It was proposed that the expression of the subgenomic flavivirus RNA (sfRNA), a viral non-coding RNA comprising vRNA 3’ NTR (and hence, predicted to associate with IGF2BP2 as ZIKV genome) alters splicing efficiency by sequestering splicing factors SF3B1 and PHAX (66). In case of DENV, NS5 was shown to associate with core components of the U5 snRNP particle and to also modulate splicing (67). If such activity is conserved for ZIKV NS5, it is tempting to speculate that mRNA splicing is regulated in ZIKV-infected cells by an IGF2BP2/NS5/sfRNA complex. We have also demonstrated that ZIKV infection specifically induces the association between IGF2BP2 and ATL2, an ER-shaping protein which was previously reported to be required for DENV and ZIKV replication (18,68). The fact that the knockdown of IGF2BP2 and ATL2 (this study and (18)) both impaired vRO biogenesis suggests that this process involves this ZIKV-specific ATL2-IGF2BP2 complex. Considering that 3’NTR is an important co-factor of VP formation (15), this complex might contribute to enriching vRNA to VPs through IGF2BP2/3’NTR interaction. It is noteworthy to mention that in DENV-infected cells, IGF2BP2 is relocalized to the replication compartment and associates with vRNA (unpublished data) but not with ATL2 (Fig. S7C-D). However, IGF2BP2 KD had very little or no effect on all tested DENV strains in contrast to ZIKV and despite our multiple attempts, we could not demonstrate evidence of a specific interaction between IGF2BP2 and DENV NS5. This suggests that i) ATL2 and NS5 are not directly involved in IGF2BP2 physical hijacking and RNA-binding activity, ii) that the physical co-opting of IGF2BP2 is not sufficient to confer the proviral activity of this host factor. However, ATL2 was reported to regulate DENV replication (18) even if it does not associate with IGF2BP2 in that infection context. Interestingly, ATL2 interactome analysis in uninfected and DENV-infected cells showed that it associates RBMX and HNRNPC, two other known “m^6^A readers” (35,69). Intriguingly, ATL2/RBMX interaction was detected only in DENV-infected cells, suggesting that it is induced by the infection and might play a similar role as the one of IGF2BP2/ATL2 in case of DENV. Finally, the interaction of IGF2BP2 with both NS5 and ATL2 was RNA-dependent (Fig. S6), which is not surprising as RNA is often a structural component of ribonucleoprotein complexes. This further supports the idea that IGF2BP2 associates with NS5 and the ER after its binding to the viral RNA.

The results described in this study allows the elaboration of a model for IGF2BP2 involvement in ZIKV life cycle (Fig. 11). In this model, very early in the life cycle, IGF2BP2 associates with vRNA and subsequently to newly synthesized NS5 (step 1). This correlates with changes in the protein composition of IGF2BP2 RNP, including induced interactions with ATL2, which target NS5/vRNA complex to the ER (step 2). Via these changes in the stoichiometry of IGF2BP2 RNP, specific endogenous IGF2BP2 mRNA ligands such as *PUM2* and *TNRC6A* mRNAs are excluded from IGF2BP2 RNP. Subsequently, this ER-bound complex induces morphological alterations of the ER membrane with the contribution of vRNA 3’ NTR and the ER-shaping activity of ATL2 to generate VPs (step 3). In addition to providing an optimal environment for vRNA synthesis, the VPs are believed to contribute to the spatial segregation of the replication complexes, assembling particles and the translation machinery, hence regulating the equilibrium between the multiple fates of vRNA. With that in mind, we cannot exclude that IGF2BP2 contributes to the coordination between replication and assembly by targeting vRNA to/through the VP pore for selective encapsidation into particles budding into the ER (step 4).

**Figure 11:**
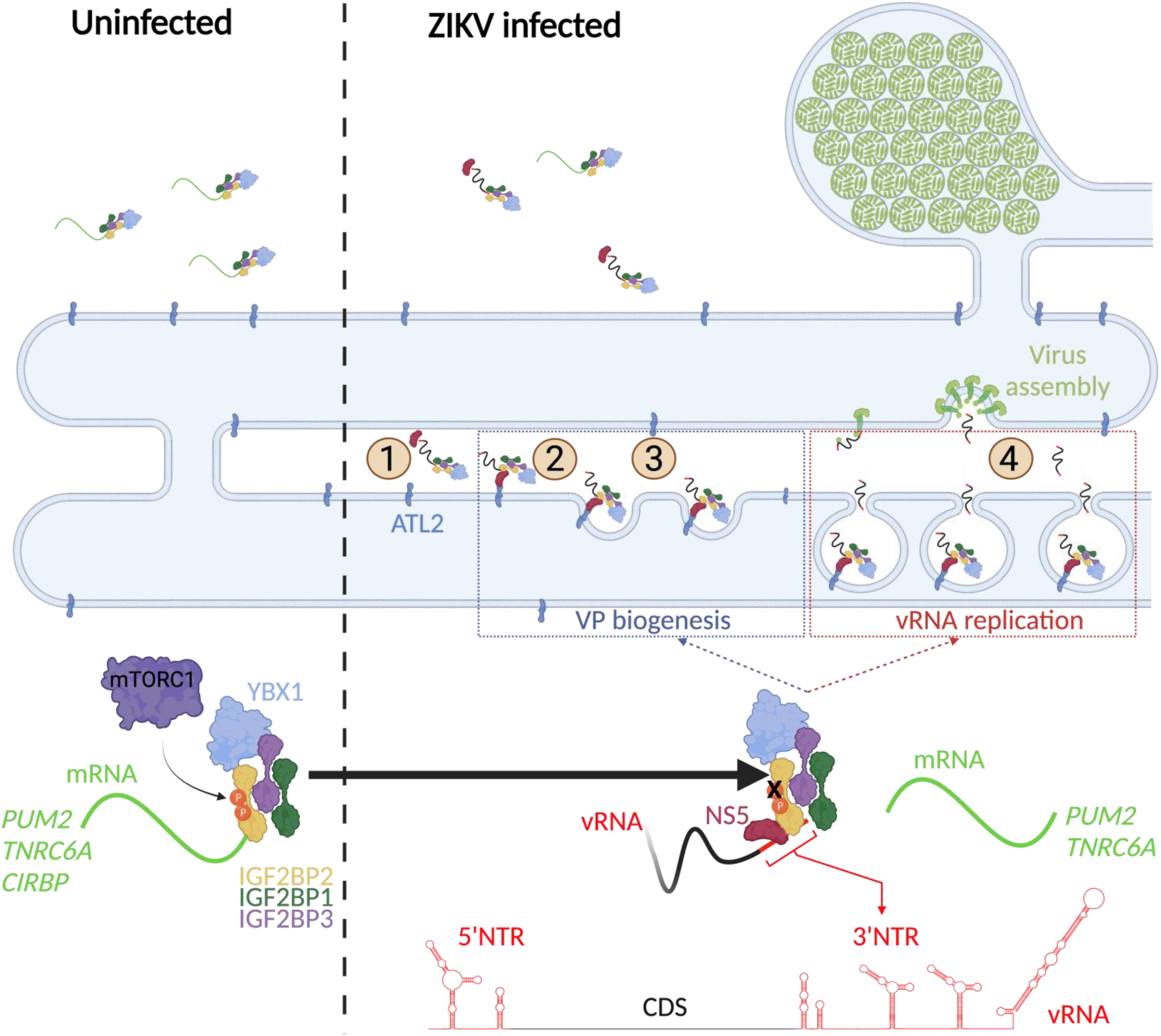
A model for IGF2BP2 involvement in ZIKV life cycle. Step 1: After NS protein synthesis early after virus entry, IGF2BP2 associates with NS5 and vRNA, thus excluding PUM2 and TNRC6A mRNA from the RNP. Step 2: The infection-induced association between IGF2BP2 RNP and ATL2 allows the targeting of vRNA/NS5 to the ER. Step 3: Viral factors and ATL2 induce the bending of the ER membrane and the formation of VPs allowing highly processive vRNA synthesis. Step 4: IGF2BP2 might be involved in the packaging of vRNA into assembling viruses by targeting the genome to the VP pore. IGF2BP2 recruitment to the replication compartment by be dependent on its mTORC1-dependent phosphorylation status.

Overall, this study highlights the physical and functional interplay between ZIKV replication machinery and IGF2BP2 RNP in infected cells. Considering the diverse roles of IGF2BP2 in mRNA metabolism, it is tantalizing to speculate that the physical and functional hijacking of IGF2BP2 RNP contributes to ZIKV pathogenesis, including developmental defects of the fetal brain.

## MATERIAL AND METHODS

### Cells, viruses and reagents

Human hepatocarcinoma Huh7.5 cells, Huh7-derived cells stably expressing the T7 RNA polymerase (Lunet-T7), HEK293T cells, Vero E6 cells, HeLa cells, NHA-htert were all cultured in DMEM (Thermo-Fisher) supplemented with 10% fetal bovine serum (FBS; Wisent), 1% non-essential amino acids (Thermo-Fisher) and 1% penicillin-streptomycin (Thermo-Fisher). The JEG-3 cell line was cultured in MEM with 10% FBS, 1% HEPES (Thermo-Fisher), 1% sodium pyruvate (Thermo-Fisher), 1% sodium bicarbonate (Thermo-Fisher) and 1% penicillin-streptomycin. Huh7.5-pWPI, Huh7.5-IGF2BP2-HA and Huh7.5-VCP-HA stable cell lines were generated by transduction of Huh7.5 with lentiviruses expressing these tagged proteins and were cultured in the presence of 1 µg/ml puromycin (Thermo-Fisher).

ZIKV H/PF/2013, ZIKV MR766 and WNV NY99 were provided by the European Virus Archive Global (EVAG). DENV1 HAWAII, DENV2 NGC, DENV3 H87, DENV4 H241 were kind gifts of Tom Hobman (University of Alberta, Canada). Virus stocks were amplified in Vero E6 cells following infection at a MOI of 0.01 for 2h. Cell supernatants were collected at 3 to 7 days post-infection, supplemented with 1% HEPES, aliquoted and stored at 80°C until use. Infectious titers were determined by plaque assays as described before (17,70,71). Plasmids containing flavivirus genomes sequences encoding Renilla luciferase (pFL-ZIKV-R2A, based on FSS13025 strain), reporter subgenomic-replicon (pFK-sgZIKV-R2A, based on H/PF/2013 strain), or replication-incompetent genomes (pFK-sgZIKV-R2A GAA) were previously reported (28,40). Plasmids were linearized using ClaI (ZIKV-R2A) or XhoI (sgZIKV-R2A and sgZIKV-R2A GAA) and subjected to *in vitro* transcription using the mMessage mMachine kit (Thermo-Fisher) with T7 RNA polymerase. DENV2 16681s, ZIKV FSS13025, ZIKV-R2A particles were produced after electroporation of VeroE6 cells with *in vitro* transcribed genomes. VeroE6 cells were resuspended in cytomix buffer (120 mM KCl, 0.15 mM CaCl_2_, 10 mM potassium phosphate buffer [pH 7.6], 25 mM HEPES [pH 7.6], 2 mM EGTA, 5 mM MgCl_2_ pH 7.6, freshly supplemented with 2 mM ATP, and 5 mM glutathione) at a density of 1.5 × 10^7^ cells per mL. 10 μg of RNA were mixed with 400 μL of cells. The cells were placed in an electroporation cuvette (0.4 cm gap width; Bio-Rad) and pulsed with a Gene Pulser Xcell Total System (Bio-Rad) at 975 μF and 270 V. Cells were seeded in a 15 cm dish. One day after electroporation, the medium was changed. Supernatants were collected, filtered through a 0.45 μm syringe filter, and supplemented with 10 mM HEPES (pH 7.5) at 3 to 7 days post-electroporation. Viruses were stored at - 80C. Stocks of SARS-CoV-2 (pre-VOC isolate Canada/QC-LSPQ-L00214517/2020 (SARS-2 LSPQ (1); GISAID : EPI_ISL_535728) were produced in VeroE6 cells. The viral titers were determined by plaque assays in Vero E6 cells.

Mouse monoclonal anti-VCP (ab11433), rabbit monoclonal anti-IGF2BP3 (ab177477) and anti-YBX1 (ab12148) were purchased from Abcam. Rabbit anti-DENV NS4B (GTX124250; cross-reactive for ZIKV), rabbit anti-ZIKV NS4B (GTX133311), rabbit anti-ZIKV NS3 (GTX133309), rabbit anti-ZIKV NS5 (GTX133312), rabbit anti-ZIKV NS4A (GTX133704), rabbit anti-DENV NS5 (GTX124253) and mouse monoclonal anti-DENV-NS3 (GTX629477; cross-reactive for ZIKV) were obtained from Genetex. Rat polyclonal antibodies targeting DENV2 16681 NS3 which are cross-reactive with ZIKV NS3 were generated at Medimabs, Montréal, Canada) and already reported (33). Mouse monoclonal anti-dsRNA (10010200) was obtained from Cedarlan. Rabbit polyclonal anti- LARP1 (A302-087A) and anti-ATL2 (A303-332A) come from Thermo Fisher Scientific. Mouse monoclonal anti-Actin (A5441) and mouse anti-HA (H3663) were purchased from Sigma-Millipore. Rabbit monoclonal anti-IGF2BP2 (14672S) and anti-IGF2BP1 (8482S) were purchased from Cell signaling.

### Plasmid design and DNA cloning

To generate tagged HA proteins IGF2BP2 and VCP expressing lentiviral constructs, PCR was performed using a plasmid VCP (wt)-EGFP gifted from Nico Dantuma (Addgene plasmid # 23971, http://n2t.net/addgene:23971; RRID:Addgene_23971) (72). Forward primers expressing HA-tag fused to VCP at the C-terminus, was used for this PCR. For IGF2BP2, we used pcDNA3-GFP-IMP2-2 (Addgene plasmid # 42175, https://www.addgene.org/42175/; RRID:Addgene_42175) as template and the HA-coding sequence was included in one of the primer to generate a C-terminally HA-tagged IGF2BP2 . PCR products were cloned into the AscI/SpeI cassette of pWPI lentiviral plasmid.

Plasmids for the recombinant expression of IGF2BP2_RRM_ and IGF2BP2_KH34_ were commercially synthesized and cloned into pET28a(+) vector (Azenta Life Science, USA). The coding sequence for each of these domains were retrieved at Uniprot (73) using the following ID Q9Y6M1. A C-terminal hexa-histidine tag was added to IGF2BP2_RRM_ and whereas a N-terminal inserted into IGF2BP2_KH34_. Additional plasmids were obtained for *in vitro* transcription of ZIKV 5’ and 3’ NTRs (Azenta Life Science, USA). cDNA sequences were obtained on GenBank (Accession ID: KJ776791.2) and inserted in pUC57 plasmid flanked by an upstream T7 promoter sequence and a XbaI cutting site downstream. A pair of guanosines were added right after the promoter sequence to increase RNA yields.

### Lentivirus production, titration and transduction

Sub-confluent HEK293T cells were cotransfected with pCMV-Gag-Pol and pMD2-VSV-G packaging plasmids and shRNA-encoding pLKO.1-puro plasmids or pWPI expressing HA-tagged proteins, using 25 kD linear polyethylenimine (Polysciences Inc.). Two- and three-days post-transfection, HEK293T supernatants were collected and filtered at 0.25 μm and stored at -80 °C. Lentiviruses were titrated in HeLa cells as previously described (26). Briefly, one day after transduction with serially diluted lentiviruses, cells are incubated with 1 μg/mL puromycin. Five days later, cell colonies were washed twice and fixed/stained with 1% crystal violet/10% ethanol for 15-30 minutes. Cells were rinsed with tap water. Colonies were counted, and titers calculated considering inoculum dilution. Huh7.5 transductions were performed using a MOI of 10 the presence of 8 µg/mL polybrene.

### Cell viability assays

MTT assays were performed to evaluate cell viability after lentiviral transduction. Huh7.5 were plated in 96-well plates (7,500 cells per well) with 8 µg/mL polybrene and with the different lentiviruses at an MOI of 10. One day posttransduction, the medium was changed and after 4 days post transduction, 20 µL of 3-(4,5-dimethylthiazol-2-yl)-2,5-diphenyltetrazolium bromide (MTT) at 5 mg/mL was added in the medium and incubated 1 to 4 hour at 37°C. Medium was removed and the MTT precipitates were dissolved with 150 µL of 2% (v/v) of 0.1 M glycin in DMSO (pH 11) per well. Absorbance at 570 nm was read with Spark® multimode microplate reader (Tecan) with the reference at 650 nm.

### Plaque assays

2.10^5^ VeroE6 cells were seeded in 24-well plates. One days after plating, virus samples were serially diluting to 10^-1^ at 10^-6^ fold in complete DMEM. 400 μl of serial dilutions were used to infect VeroE6 cells in duplicates (200 μl of dilution/well). 2h post-infection, the inoculum was removed and replaced for serum-free MEM (life Technologies) containing 1.5% carboxymethylcellulose (Millipore-Sigma) for ZIKV, DENV and WNV. For SARS-CoV-2 infected cells were incubated in MEM-0.8% carboxymethylcellulose. After 7 days for DENV2 16681s, 6 days for DENV1 HAWAII, 5 days DENV2 NGC and DENV3 H84, 4 days for ZIKV H/PF/2013, ZIKV MR766, ZIKV FSS13025 and DENV4 H241 and 3 for days WNV NY99 and SARS-CoV-2, titration are fixed during 2 hours in 2.5% formaldehyde. After thoroughly rinsing with tap water, cells were stained with 1% crystal violet/10% ethanol for 15-30 minutes. Stained cells were washed with tap water, plaques were counted, and titers of infectious viruses were calculated in PFU/mL.

### Renilla luciferase assay

10^5^ Huh7.5 cells were transduced and seeded in 12-well plates. One day post-transduction, we changed the medium, and the day after, we infected them with ZIKV-R2A reporter viruses at a MOI of approximately 0.0001. 2 days post-infection, cells were lysed in 200µl of luciferase lysis buffer (1% Triton-X-100; 25mM Glycyl-Glycine, pH7.8; 15mM MgSO_4_; 4 mM EGTA; 1 mM DTT added directly prior to use). 30 µl of lysates were plated in a 96-well white plate and luminescence was read with a Spark multimode microplate reader (Tecan) after injection of 150 µl of luciferase buffer (25mM Glycyl-Glycine pH7.8; 15mM KPO_4_ buffer pH7.8; 15mM MgSO_4_; 4mM EGTA; 1mM coelenterazine added directly prior to use). All values were background-subtracted and normalized to the control shNT-transduced cells.

### Immunofluorescence assays

Huh 7.5 cells were grown in 24-well plates containing sterile coverslips. 1-day post-seeding, cells were infected H/PF/2013 with an MOI of 5-10. Two-day post-infection, cells were washed three times with PBS before fixation, with 4% PFA for 20 minutes. Then, cells were washed 3 times with PBS before storing at 4 °C. Cells were permeabilized with PBS 1X-0.2% Triton X-100 for 15 minutes at room temperature, and subsequently blocked for 1h with PBS supplemented with 5% bovine serum albumin (BSA) and 10% goat serum. Cells were incubated with primary antibodies for two hours protected from light at room temperature. Coverslips were washed three times with PBS before incubating for 1h with Alexa Fluor (488, 568 or 647)-conjugated secondary antibodies, in the dark at room temperature. Coverslips were then washed three times for 10 minutes with PBS and incubated for 15 minutes with 4’, 6’-diamidino-2-phenylindole (DAPI; Life Technologies) diluted 1:10000 in PBS. Finally, coverslips were washed three times with PBS and once with distilled water before mounting on slides with FluoromountG (Southern Biotechnology Associates). Imaging was carried out with a LSM780 confocal microscope (Carl Zeiss Microimaging) at the Confocal Microscopy Core Facility of the INRS-Centre Armand-Frappier Santé Biotechnologie and subsequently processed with the Fiji software. For the colocalization analysis, the Manders’ coefficients were measured using the JacoP plugin in Fiji. In this study, the Manders’ coefficient represents the fraction of NS3 or dsRNA signals overlapping with the signals of the indicated cellular RNA-binding proteins.

### Co-immunoprecipitation assays

For anti-HA immunoprecipitation, cells stably expressing IGF2BP2-HA protein, or VCP-HA protein, or transduced with the control lentiviruses (pWPI) were infected with ZIKV H/PF/2013 at a MOI of 10. Two days post-infection, cells were washed twice with PBS, collected and lysed for 20 minutes on ice in a buffer containing 0.5% NP-40, 150 mM NaCl, 50 mM TrisCl pH 8.0 and EDTA-free protease inhibitors (Roche). The cell lysates were centrifuged at 13,000 rpm for 15 minutes at 4 °C and supernatants were collected and stored at -80C. Bradford assays were carried out to quantify protein concentration, before performing immunoprecipitation with the equal quantities of protein for each condition (350-800 μg of proteins) in 1-1.5 mL total volume. For Figure S6, prior to the immunoprecipitation step, cell extracts were treated with 20µg/mL RNase A for 20 minutes at room temperature. The HA-immunoprecipitation was performed by incubating the lysated with 50 μL of a 50/50 slurry of mouse monoclonal anti-HA coupled to agarose beads (Millipore-Sigma) for 3 h at 4 °C on a rotating wheel. The resin was washed 4 times with lysis buffer. The last wash was performed in new low binding microtubes. For immunoblotting, complexes were eluted from the resin with standard SDS-PAGE loading buffer. for RNA extraction, beads were resuspended in 250 μl lysis buffer and 1ml Trizol-LS (Life Technologies) was added. The samples were stored at -80 °C before western blotting. The quantification of western blots signals was performed with the ImageLab software (Bio-Rad).

For the mass spectrometry experiment, the immunoprecipitated samples were produced with the same protocol, but the final wash steps are different. After lysate incubation with the antibodies, the first 3 washes were done with 1mL of lysis buffer (10 minutes each) at 4°C on a rotating wheel and spun down 1 min at 10,000 rpm. The last quick wash was done with detergent-free washing buffer (150 mM NaCl; 50 mM TrisCl pH 8.0) and spun down for 1 min at 10,000 rpm. After the removal of buffer excess, and using a cut 200 μL tips, beads were transferred into new low-binding tubes. We performed 3 quick washes with 1mL of washing buffer. After removal of the buffer, the samples were snap-frozen in ice dry and stored at -80 °C until use.

### RT-qPCR

Total RNA was extracted from cells using the RNeasy Mini kit (Qiagen). Co-immunoprecipitated RNA was extracted from immunoprecipitated samples using Trizol-LS according to the manufacturer’s instructions. RNAs were subjected to reverse transcription using the Invitrogen SuperScript IV VILO Master Mix RT kit (Life Technologies). Real-time PCR was performed using the Applied Biosystems SYBR green Master mix (Life Technologies) and a LightCycler 96 (Roche) for the detection. The following primer pairs were used: ZIKV H/PF/2013 vRNA: 5′-AGATGAACTGATGGCCGGGC-3′ and 5′-AGGTCCCTTCTGTGGAAATA-3′; *GAPDH* 5′-GAAGGTGAAGGTCGGAGTC-3′ and 5′-GAAGATGGTGATGGGATTTC-3′; *IGF2BP2* 5’-CGGGGAAGAGACGGATGATG-3’ and 5’-CGCAGCGGGAAATCAATCTG-3’; *PUM2* 5’-TTTGCGCAAATACACATACGGG-3’ and 5’-GGTCCTCCAATAGGTCCTAGGT-3’; *CIRBP* 5’-GACCACGAGCCATGAGTTTTC-3’ and 5’-CTCAGAGAAGTGAGTGGGGC-3’; *TNRC6A* 5’-ACTAACTGTGGAGACCTTCACG-3’ and 5’-GTTAATGGGAGATGGGCTGCTA-3’. Relative abundance of RNAs was calculated with the ΔΔCt method using *GAPDH* mRNA as a reference RNA.

### RNA fluorescence in situ hybridization

Coverslip were coated with collagen I (Corning) for 30 minutes and washed 3 times with PBS. 7×10^4^ Huh 7.5 cells/well were seeded in on 24-well plates and infected one day later with ZIKV H/PF/2013 with a MOI of 10. Two-day post-infection, cells were washed three times with PBS before fixation with 4% PFA for 20 minutes. Then, cells were washed 3 times with PBS before storing at 4 °C. FISH was performed using ViewRNA ISH Cell Assay kit (Thermo-Fisher) according to the manufacturer’s instructions. Briefly, cells were permeabilized with the kit QS detergent for 5 minutes, followed by 3 PBS washes and kit protease QS treatment. The protease QS was diluted to 1:10,000 and incubated 10 minutes at room temperature. The probes for ZIKV RNA recognition (VF4-20142; Thermo-Fisher) were diluted 1:100 with the kit dilution buffer. Probe hybridization was performed at 40 °C for 3h. Subsequently, the pre-amplifying, amplifying and fluorophore labeling steps were performed, each at 40 °C for 30 minutes. We wash the cells like it is mentioned in the protocol. Coverslips were then immunostained with anti-IGF2BP2 antibodies and AlexaFluor488 secondary antibodies followed by DAPI staining using a standard immunofluorescence protocol. For the colocalization analysis, the Manders’ coefficient representing the fraction of vRNA signal overlapping with IGF2BP2 signal was measured using the JacoP plugin in Fiji.

### IGF2BP2 protein production for *in vitro* assays

For the expression of IGF2BP2_RRM_, plasmid was transformed into Rosetta^TM^ DE3 competent cells (Sigma Aldrich) by heat shock. A single colony was selected and grown in 50 mL of LB media for 16-18 hours at 37 °C and 220 rpm, and then transferred to a secondary culture of 500 mL of LB. Secondary cultures were incubated at 37°C and 220rpm before induction with 1 mM IPTG when the cultures reached OD_600_ reached 3.0. Following induction, cultures were incubated at 20°C and 220 rpm for 16-18 hours. Cells were centrifuged and resuspended in lysis buffer (50 mM phosphate pH 7.0, 500 mM NaCl, 10 mM imidazole). Next, we incubated the cells for 20 minutes at 4 °C following the addition of 70 μM lysozyme, 1 μM PMSF, 80 mU/mL DNase, and 3 μM sodium deoxycholate. The cells were then lysed via sonication. Lysate was then centrifuged and passed over a HisTrap^TM^ HP (Global Life Science Solutions, USA) column. The column was washed with lysis buffer containing 30 mM imidazole, then eluted with a gradient elution by mixing lysis and elution buffers (50 mM phosphate pH 7.0, 500 mM NaCl, and 250 mM imidazole). The purity and homogeneity of selected samples were assessed by SDS-PAGE immediately after purification. Selected fractions were concentrated in Vivaspin® 20 10kDa MWCO centrifugal concentrators and applied to a Mono Q^TM^ 10/100 GL column (Global Life Science Solutions, USA) equilibrated with storage buffer (50 mM phosphate pH 7.0, 150 mM NaCl) to remove any residual nucleic acid. Eluted fractions were loaded in SDS-PAGE and homogenous samples were again pooled and concentrated, then passed through a Superdex^TM^ 75 Increase 10/300 GL column (Global Life Science Solutions, USA) equilibrated with storage buffer as a final purification step. Samples were once again checked with SDS-PAGE, and homogenous samples were stored at 4°C until needed.

Expression and purification of IGF2BP2KH3-4 was performed as described previously (74). Briefly, vectors were transformed and grown to an OD of 0.6, then induced with 1 mM IPTG. Cells were centrifuged and resuspended in lysis buffer (50 mM phosphate pH 7.0, 1.5 M NaCl, 10 mM imidazole) and then lysed through sonication. Purification was performed using nickel affinity chromatography, with eluted fractions being checked using SDS-PAGE for homogeneity. Samples were concentrated and passed over a Superdex^TM^ 75 Increase 10/300 GL column as a final purification step.

### *In vitro* transcription and RNA labelling

Plasmids containing cDNA for ZIKV 5’ and 3’ NTRs were transformed and cultured in *E. coli* NEBα (NEB, Canada) competent cells and further recovered using GeneJET Plasmid Maxiprep Kit (ThermoFisher Scientific, Canada) as per manufacturer’s protocol, followed by a 4-hour linearization with XbaI endonuclease (NEB, Canada) at 37 °C. ZIKV NTRs RNAs were synthesized by *in vitro* transcription using an in-house purified T7 polymerase, as previously described (75,76). Subsequently, RNAs were purified on a Superdex^TM^ 200 10/300 GL column (Global Life Science Solutions, USA) equilibrated with RNA buffer (10 mM Bis-tris pH 6.0, 100 mM NaCl, 15 mM KCl, 5 mM MgCl_2_, 5% glycerol).

Pure and homogeneous RNAs were concentrated to ∼ 100 μM through ethanol precipitation and then fluorescently labelled at the 5’ end. The labelling reaction was then prepared from 30 μL of concentrated RNA, 1.25 mg of 1-ethyl-3-(3-dimethylamino) propyl carbodiimide hydrochloride (EDC), and 5 mg/mL of Alexa Fluor 488 dye (Thermo Fisher Scientific, USA) dissolved in DMSO. Samples were thoroughly mixed until contents were entirely dissolved before adding 20 μL of 0.1 M imidazole, pH 6. Afterward, we incubated the samples at room temperature for 3 hours before being further purified on a Superdex^TM^ 200 10/300 GL column. RNA was checked for degradation using agarose gel electrophoresis and for labelling efficiency using microscale thermophoresis (MST).

### Microscale thermophoresis-based *In vitro* RNA binding assays

MST experiments were conducted at room temperature using Nanotemper Technologies Monolith® NT.115 instrument to measure binding affinity. Pure fluorescently labelled RNAs were used as targets with a fixed concentration of 250 nM for both ZIKV 5’ and 3’ NTRs. The proteins were set as the ligand and diluted in a 2-fold serial dilution with concentrations ranging from 65.00 to 1.98 x 10^4^ μM for IGF2BP2_KH34_ and 10.00 to 6.10 x 10^5^ μM for IGF2BP2_RRM_. Reactions were prepared using MST buffer (50 mM phosphate pH 7.0, 150 mM NaCl, and 0.05% Tween 20), incubated at room temperature for 20 minutes and subsequently loaded into standard capillaries. The binding affinity experiments were performed using medium MST-Power, 80% excitation-power, with data collection on the cold region at 0 s and the hot region at 5 s. Three independent replicates were collected and analyzed using MO.Affinity Analysis software v2.1.3, in which K_d_ fitting models were obtained and plotted using GraphPad Prism 9 software.

### Affinity-purification and quantitative LC-MS/MS proteomics

For the determination of IGFBP2 interactome, five independent affinity purifications using an anti-HA conjugated agarose beads (Millipore-Sigma) were performed for each experimental condition. Confluent monolayers of Huh7.5 cells constitutively expressing empty ctrls (NT) or HA-tagged IGFBP2 were mock-infected or infected with DENV or ZIKV at an MOI of 5 and 10, respectively. Cells were lysed in Lysis Buffer (50 mM Tris pH 7.6, 150 mM NaCl, 0.5% NP-40) containing protease and phosphatase inhibitors (cOmplete and PhosStop, Roche), and processed as previously described (PMID: 30177828). Bound proteins (IP) or 50 µg of normalized whole cell lysates (Input) were denatured by incubation in 40 µl U/T buffer (8 M Urea, 6M Thiurea, 100 mM Tris-HCl pH 8.5), and reduction and alkylation carried out with 10 mM DTT and 55 mM iodoacetamide in 50 mM ABC buffer (50 mM NH_4_HCO_3_ in water pH 8.0), respectively. After digestion with 1 µg LysC (WAKO Chemicals USA) at room temperature for 3 h, the suspension was diluted in ABC buffer, and the protein solution was digested with trypsin (Promega) overnight at room temperature. Peptides were purified on stage tips with three C18 Empore filter discs (3M) and analyzed by liquid chromatography coupled to mass spectrometry as previously described (77).

Samples were analysed on a nanoElute (plug-in v.1.1.0.27; Bruker) coupled to a trapped ion mobility spectrometry quadrupole time of flight (timsTOF Pro) (Bruker) equipped with a CaptiveSpray source. Peptides were injected into a Trap cartridge (5 mm × 300 μm, 5 μm C18; Thermo Fisher Scientific) and next separated on a 25 cm × 75 μm analytical column, 1.6 μm C18 beads with a packed emitter tip (IonOpticks). The column temperature was maintained at 50 °C using an integrated column oven (Sonation GmbH). The column was equilibrated using 4 column volumes before loading samples in 100% buffer A (99.9% Milli-Q water, 0.1% formic acid (FA)). Samples were separated at 400 nl min−1 using a linear gradient from 2 to 17% buffer B (99.9% ACN, 0.1% FA) over 60 min before ramping up to 25% (30 min), 37% (10 min) and 95% of buffer B (10 min) and sustained for 10 min (total separation method time, 120 min). The timsTOF Pro was operated in parallel accumulation-serial fragmentation (PASEF) mode using Compass Hystar v.5.0.36.0. Settings were as follows: mass range 100– 1700 m/z, 1/K0 start 0.6 V⋅s/cm2End 1.6 V⋅s/cm2; ramp time 110.1 ms; lock duty cycle to 100%; capillary voltage 1,600 V; dry gas 3 l min−1; dry temperature 180 °C. The PASEF settings were: 10 tandem mass spectrometry (MS) scans (total cycle time, 1.27 s); charge range 0–5; active exclusion for 0.4 min; scheduling target intensity 10,000; intensity threshold 2,500; collision-induced dissociation energy 42 eV.

### Raw data processing and analysis

Raw MS data were processed with the MaxQuant software v.1.6.17 using the built-in Andromeda search engine to search against the human proteome (UniprotKB, release 2019_10) containing forward and reverse sequences concatenated with the DENV-2 16681 strain (UniprotKB #P29990) and ZIKV H/PF/2013 strain (UniprotKB #KU955593) with the individual viral open reading frames manually annotated, and the label-free quantitation algorithm (78). Additionally, the intensity-based absolute quantification (iBAQ) algorithm and match between runs option were used. In MaxQuant, carbamidomethylation was set as fixed and methionine oxidation and N-acetylation as variable modifications. Search peptide tolerance was set at 70 p.p.m. and the main search was set at 30 p.p.m. (other settings left as default). Experiment type was set as TIMS-DDA with no modification to the default settings. Search results were filtered with a false discovery rate of 0.01 for peptide and protein identification. The Perseus software v.1.6.15 was used to process the data further. Protein tables were filtered to eliminate the identifications from the reverse database and common contaminants. When analysing the MS data, only proteins identified on the basis of at least one peptide and a minimum of three quantitation events in at least one experimental group were considered. The iBAQ protein intensity values were normalized against the median intensity of each sample (using only peptides with recorded intensity values across all samples and biological replicates) and log-transformed; missing values were filled by imputation with random numbers drawn from a normal distribution calculated for each sample. PCA analysis was used to quality control variance across biological replicates, and remove outliers (replicate 1 and 5 of IGFBP2 pull-downs were removed from all the matrix).

Significant interactors were determined by multiple ANOVA T-tests with permutation-based false discovery rate statistics. We performed 250 permutations, and the FDR threshold was set at 0.05. The parameter S0 was set at 1 to separate background from specifically enriched interactors. Unsupervised hierarchical clustering of proteins was performed on logarithmized and z-scored intensities of ANOVA significant interactors. Five unique clusters of proteins differentially regulated upon viral infection were identified (Fig. S7). Results were plotted as scatter plot and heat map using Perseus(79) or Adobe Illustrator.

### Accession numbers & Data Availability

UniprotKB accession codes of all protein groups and proteins identified by Mass Spectrometry are provided in each respective Supplementary Table and were extracted from UniprotKB (Human; release 2019_10). The mass spectrometry proteomics data have been deposited to the ProteomeXchange Consortium (http://proteomecentral.proteomexchange.org) via the PRIDE partner repository with the dataset identifier PXD052835.

### Viral replication organelle induction and imaging

Lunet T7 cells expressing a cytosolic T7 polymerase, were seeded into 6-well plates and transduced with lentivirus (MOI 5) containing shRNA sequences targeting either IGF2BP2 or nontarget (NT). After 2 days, transduced cells were then seeded into 24-well plates (30,000 cells/well). One day later, cells were transfected with the pIRO-Z system as described in (Goellner et. al. 2020 and Cerikan et. al. 2020), using Mirus TransIT transfection reagent. After 18 hours, cells were fixed with 1.25% glutaraldehyde in 0.2 M HEPES buffer (for EM imaging) or processed for Immunofluorescence, western blot, or RT-qPCR as described above.

To prepare cells for EM imaging, glutaraldehyde fixed samples were washed three times with 0.2 M HEPES. Next, samples were incubated with 1% osmium tetroxide/0.1 M Cacodylate for 1 hour and washed three times with HPLC grade water. Samples were then incubated with 2% uranyl acetate for 30 minutes at 60℃ followed by washing with HPLC-grade water three times. Progressive dehydration was performed with increasing concentrations of ethanol (40% to 100%). Cells were embedded into an Eponate 12 Resin (Ted Pella inc) and left for two days at 60°C to achieve complete polymerization. Embedded samples were sectioned into 70-nm-thick slices using a Leica EM UC6 microtome and a diamond knife (Diatome). Sections were counterstained with 4% uranyl acetate for 5 minutes and 2% lead citrate for 2 minutes.

Sections were then mounted on EM grids for imaging. Electron microscopy images were obtained on a JEOL JEM-1400 120 kV LaB6 transmission electron microscope with a Gatan US1000 CCD camera at 8000X magnification. Quantification was performed by systematically surveying cells and evaluating the presence of VPs. Only cells with >2 VPs were considered as positive. For each condition, >50 cells were surveyed over 4 biological replicas. All observed VPs were imaged, and VP diameters were determined using ImageJ by measuring the distance across two axes and averaging.

### Bio-informatic analyses

The gene ontology analysis was performed with ShinyGO 0.77 software (http://bioinformatics.sdstate.edu/go/). The IGF2BP2 interactomic tree was generated with STRING database using the online resource at https://string-db.org/.

### Statistical analysis

Statistical analyses were performed with GraphPad Prism 9 software. In the figures 1, 9 and S7, the statistical significance was evaluated by a one-way ANOVA test. For the figures 2 to 9, 11, S6 and S7, the statistical significance was determined using unpaired *t*-tests. p value below 0.05 were considered significant: ****: p<0.0001; ***: p< 0.001; **: p < 0.01; * p< 0.05.

## ACKNOWLEDGEMENTS

We thank Dr. Alessia Ruggieri (University of Heidelberg) and Dr. Mirko Cortese (Telethon Institute of Genetics and Medicine) for excellent scientific discussion about this project. We are grateful to Jessy Tremblay and Arnaldo Nakamura at the Confocal Microscopy and Flow Cytometry Facility and the Electron Microscopy Facility of INRS-Centre Armand-Frappier, respectively for excellent technical assistance. The authors wish to acknowledge the support of the Emory University Robert P. Apkarian Integrated Electron Microscopy Core Facility (RRID: SCR_023537) for help with preparing EM samples and with establishing imaging workflows. We thank Dr. Pei-Yong Shi, the World Reference Center for Emerging Viruses and Arboviruses (WRCEVA) and Dr. Ralf Bartenschlager (University of Heidelberg) for providing the ZIKV and DENV reverse genetics systems, and Dr. Daniel Lamarre, Dr. Frédérick Antoine Mallette (University of Montréal), Dr. Tom Hobman (University of Alberta), Dr. Patrick Labonté (Institut National de la Recherche Scientifique) and Dr. Anil Kumar (University of Saskatchewan) for generously providing us with shRNA-expressing lentiviral plasmids and cell lines. We are grateful to the European Virus Archive Global (EVAg), Dr. Xavier de Lamballerie (Emergence des Pathologies Virales, Aix-Marseille University, France) and Robin Gafur (Animal and Plant Health Agency, Addlestone, UK) for providing ZIKV H/PF/2013 and WNV99 original stocks. We thank the Laboratoire de Santé Publique du Québec and Philippe Dufresne for providing the SARS-CoV-2 isolate, and Dr. Alain Lamarre and Mrs Tania Charpentier for help with the amplification of this virus. C.M and A.A.S received PhD fellowships from the Armand-Frappier Foundation and the Center of Excellence in Research on Orphan Diseases-Courtois Foundation (CERMO-FC). A.A.S is receiving a PhD fellowship from the Fonds de Recherche du Québec-Nature et Technologies (FRQNT). H.S.P is receiving a CREATE Postdoctoral Fellowship award from the Natural Sciences and Engineering Research Council of Canada (NSERC). T.R.P is supported by a NSERC Discovery grant (RGPIN-2022-03391) and the Canada Foundation for Innovation (CFI-37155 and CFI-41008), and acknowledges the Canada Research Chair program. L.C.C is receiving a senior research scholar salary support from Fonds de Recherche du Québec-Santé (FRQS). Work in P.S. laboratory is funded by the Free and Hanseatic City of Hamburg, the German Federal Ministry of Education and Research (VirMScan) and the German Research Foundation (SC 314/1-2). This project was supported by a Discovery grant from NSERC (RGPIN-2016-05584) and an Early Career Investigator grant from FRQNT (2018-NC-205593), awarded to L.C.C.

---

**Figure S1:**
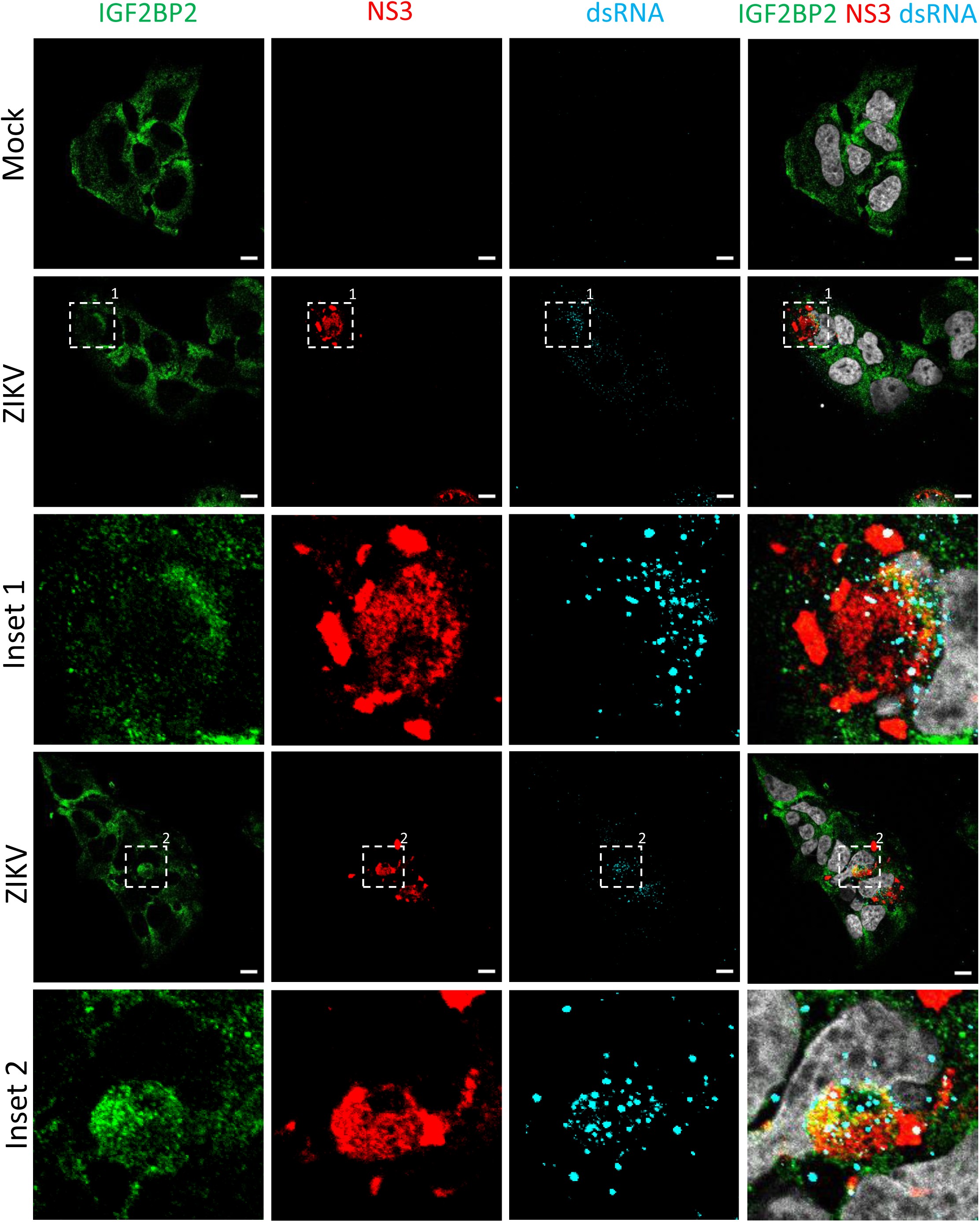
IGF2BP2 relocalizes to the viral replication compartment in ZIKV-infected cells at one day post-infection. Huh7.5 cells were infected with ZIKV H/PF/2013 with a MOI of 10 or left uninfected. At one day post-infection, cells were fixed, immunolabelled for the indicated factors, and imaged by confocal microscopy. Scale bar=10 µm. The white squares indicate the magnified areas in the insets.

**Figure S2:**
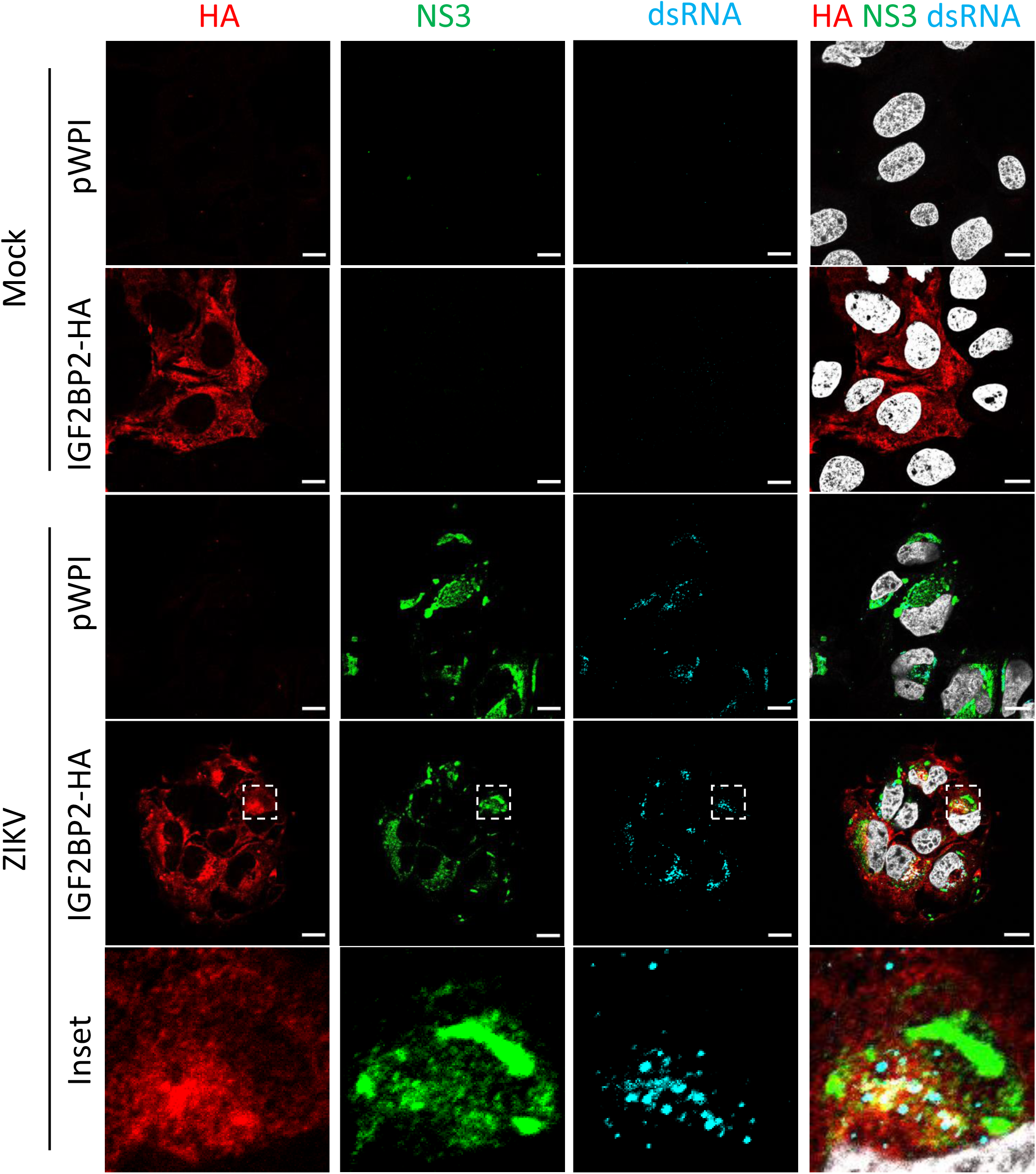
HA-tagged IGF2BP2 relocalizes to the viral replication compartment in ZIKV-infected cells, as endogenous IGF2BP2. Huh7.5 cells stably overexpressing IGF2BP2-HA or control transduced cells (pWPI) were infected with ZIKV H/PF/2013 with a MOI of 10 or left uninfected. Two days post-infection, cells were fixed, immunolabelled for the indicated factors, and imaged by confocal microscopy. Scale bar=10 µm. The white squares indicate the magnified areas in the insets.

**Figure S3:**
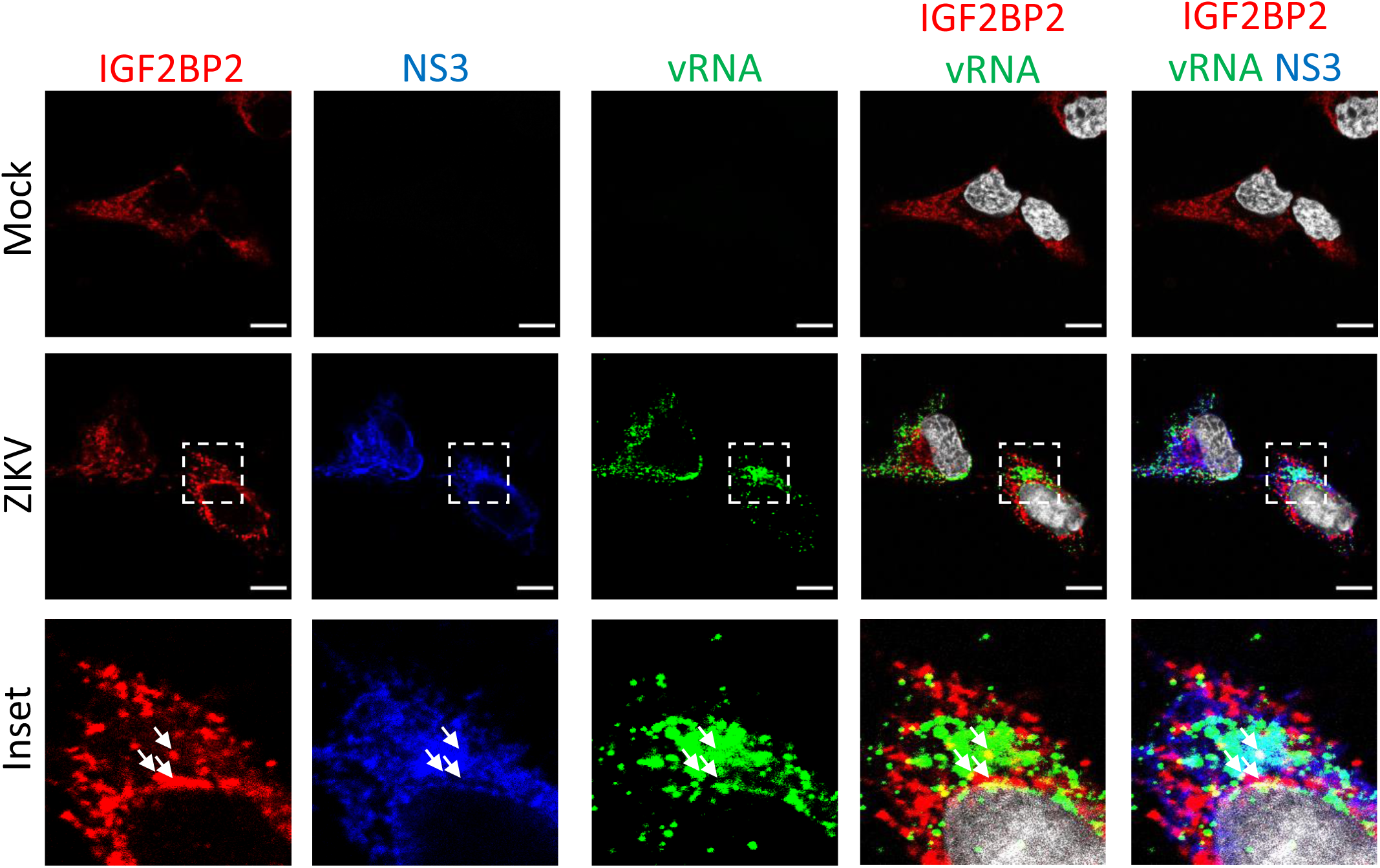
IGF2BP2 partially co-localizes with NS3 and vRNA in ZIKV infected cells. FISH with IGF2BP2 and NS3 co-immunostaining was performed in Huh7.5 cells after 2 days post infection with ZIKV at an MOI of 10. White arrows show the triple colocalization. Scale bar=10 µm. The white squares indicate the magnified areas in the insets.

**Figure S4:**
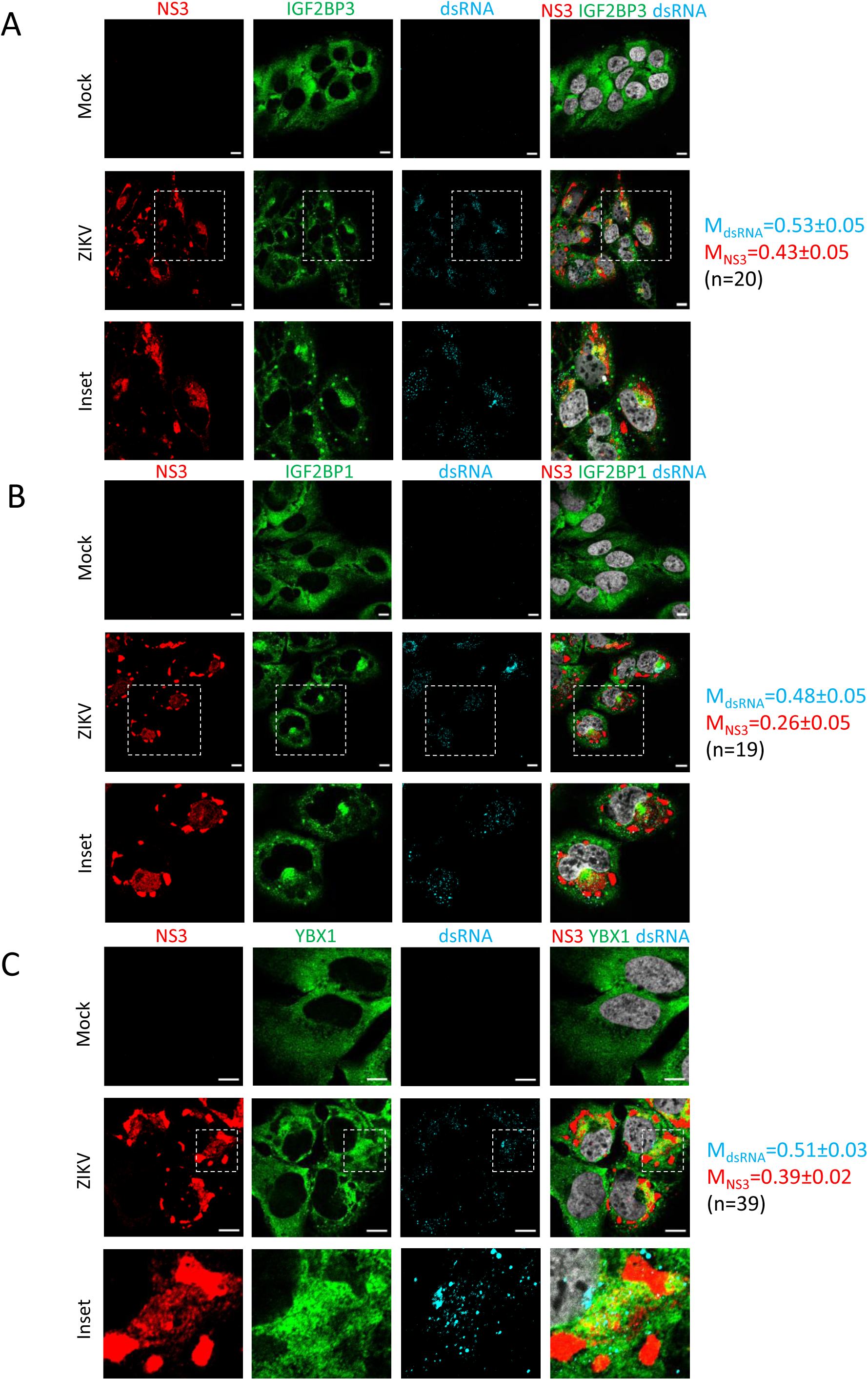
IGF2BP1, IGF2BP3 and YBX1 relocalize to the viral replication compartment in ZIKV-infected cells. Huh7.5 cells were infected with ZIKV H/PF/2013 with a MOI of 10 or left uninfected. Two days post-infection, cells were fixed, immunolabelled for the indicated factors, and imaged by confocal microscopy. Scale bar=10 µm. The Manders’ coefficients (mean ± SEM) representing the fraction of dsRNA (cyan) and NS3 (red) signals overlapping with IGF2BP3 (A), IGF2BP1 (B) or YBX1 (C) signals are shown (n=number of cells). The white squares indicate the magnified areas in the insets.

**Figure S5:**
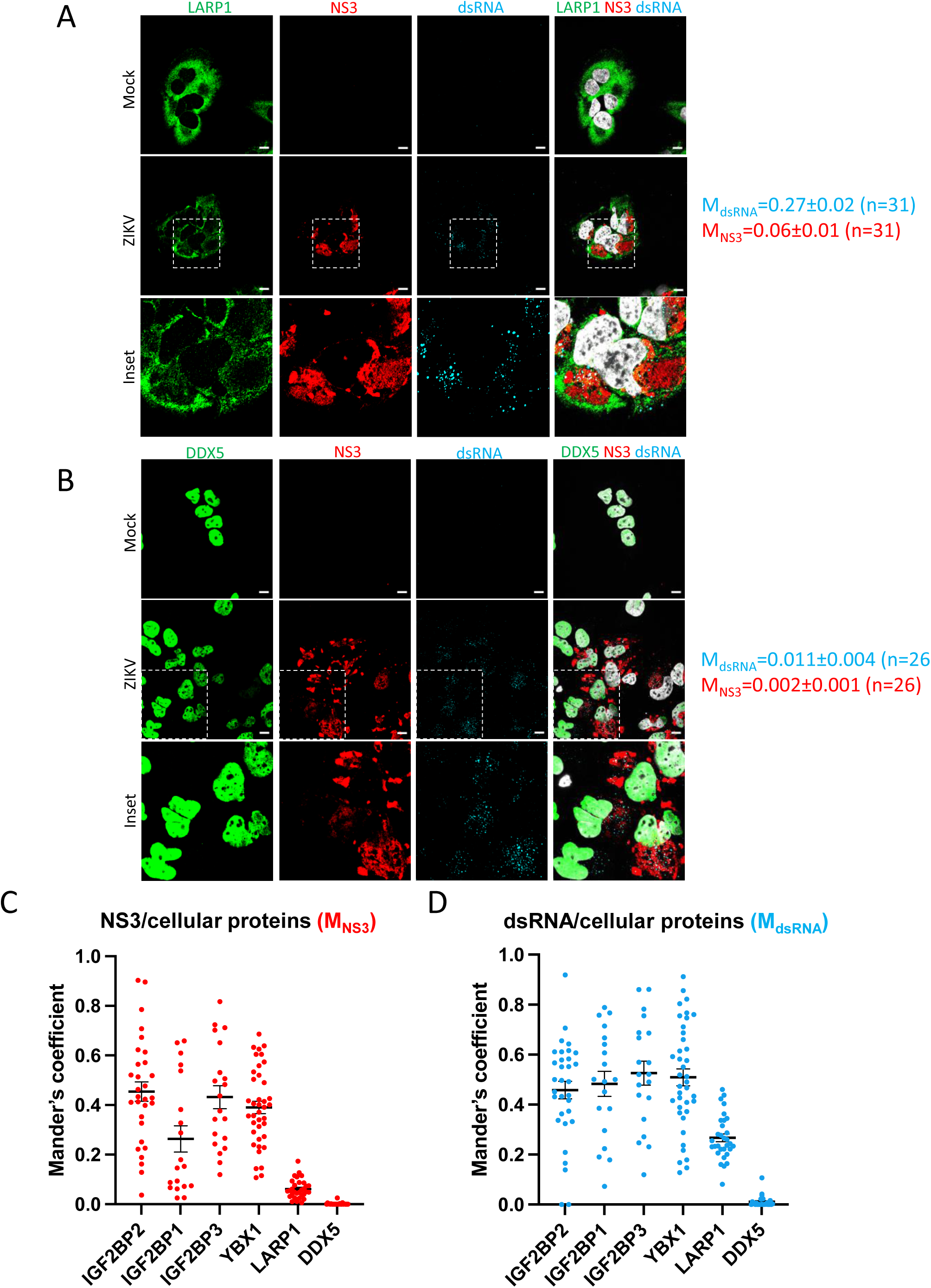
LARP1 and DDX5 do not relocalize to the viral replication compartment in ZIKV-infected cells. (A-B) Huh7.5 cells were infected with ZIKV H/PF/2013 with a MOI of 10 or left uninfected. Two days post-infection, cells were fixed, immunolabelled for the indicated factors, and imaged by confocal microscopy. Scale bar=10 µm. The Manders’ coefficients (mean ± SEM) representing the fraction of dsRNA (cyan) and NS3 (red) signals overlapping with LARP1 (A) or DDX5 (B) signals are shown (n=number of cells). The white squares indicate the magnified areas in the insets. (C-D) The Manders’ coefficients per cell determined from Figures 4B, S4A-C and S5A-B are plotted. Means ± SEM are shown in black.

**Figure S6:**
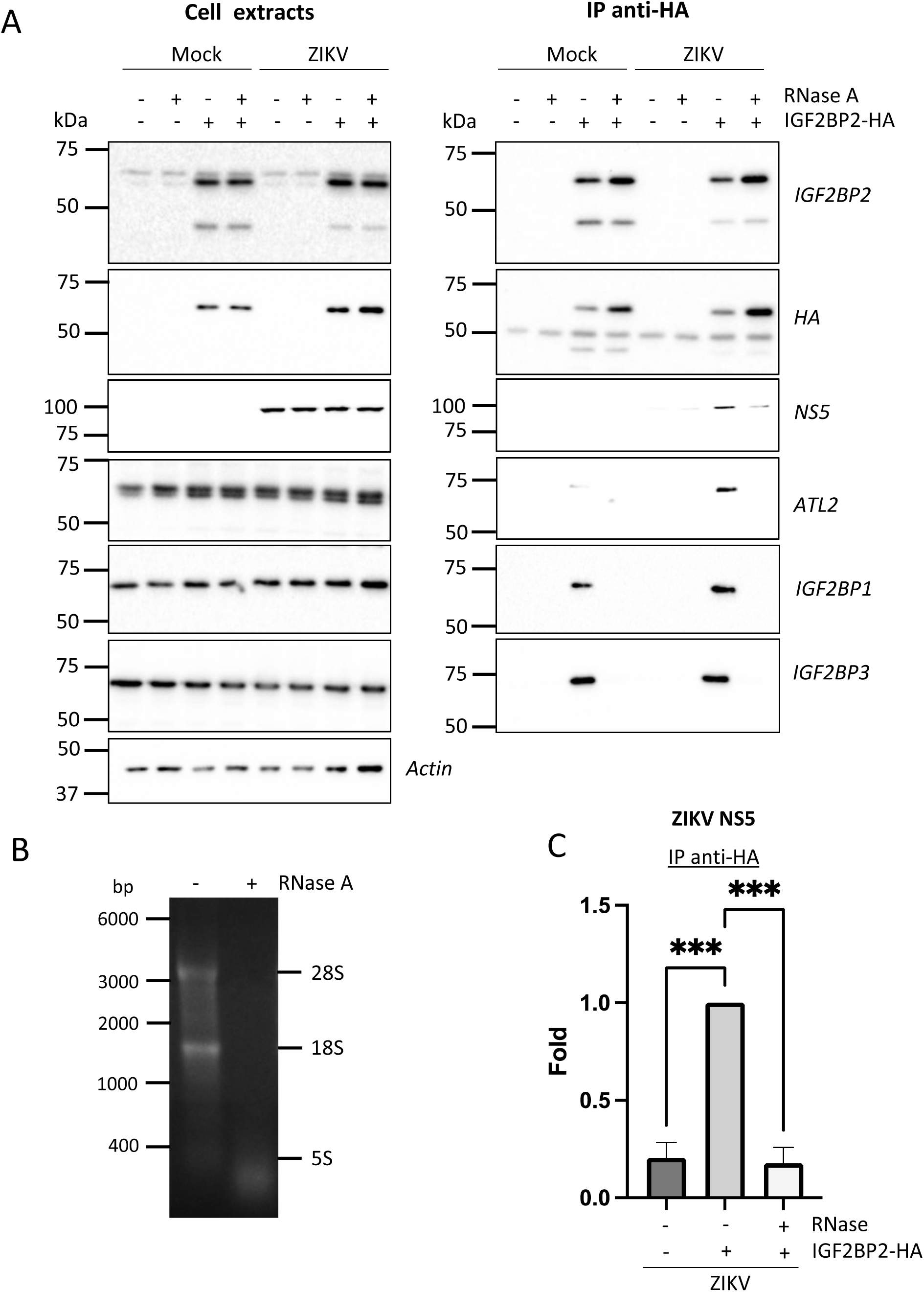
The association between IGF2BP2 and ZIKV NS5 is RNA-dependent in infected cells. (A) Huh7.5 cells stably expressing IGF2BP2-HA (+) and control cells (-) were infected with ZIKV H/PF/2013 at a MOI of 10 or left uninfected. Two days later, cell extracts were prepared and subjected to RNase A treatment (+) or not (-) before anti-HA immunoprecipitations. The resulting complexes were analyzed by western blotting for their abundance in the indicated proteins. (B) The RNA content in cell extracts was analyzed on an agarose gel for controlling the efficiency of the RNase A treatment. (C) ZIKV NS5 levels in the IP samples were quantified and means of protein signals (normalized to IGF2BP2) ± SEM based on three independent experiments are shown. ***: p<0.001 (unpaired t-test).

**Figure S7:**
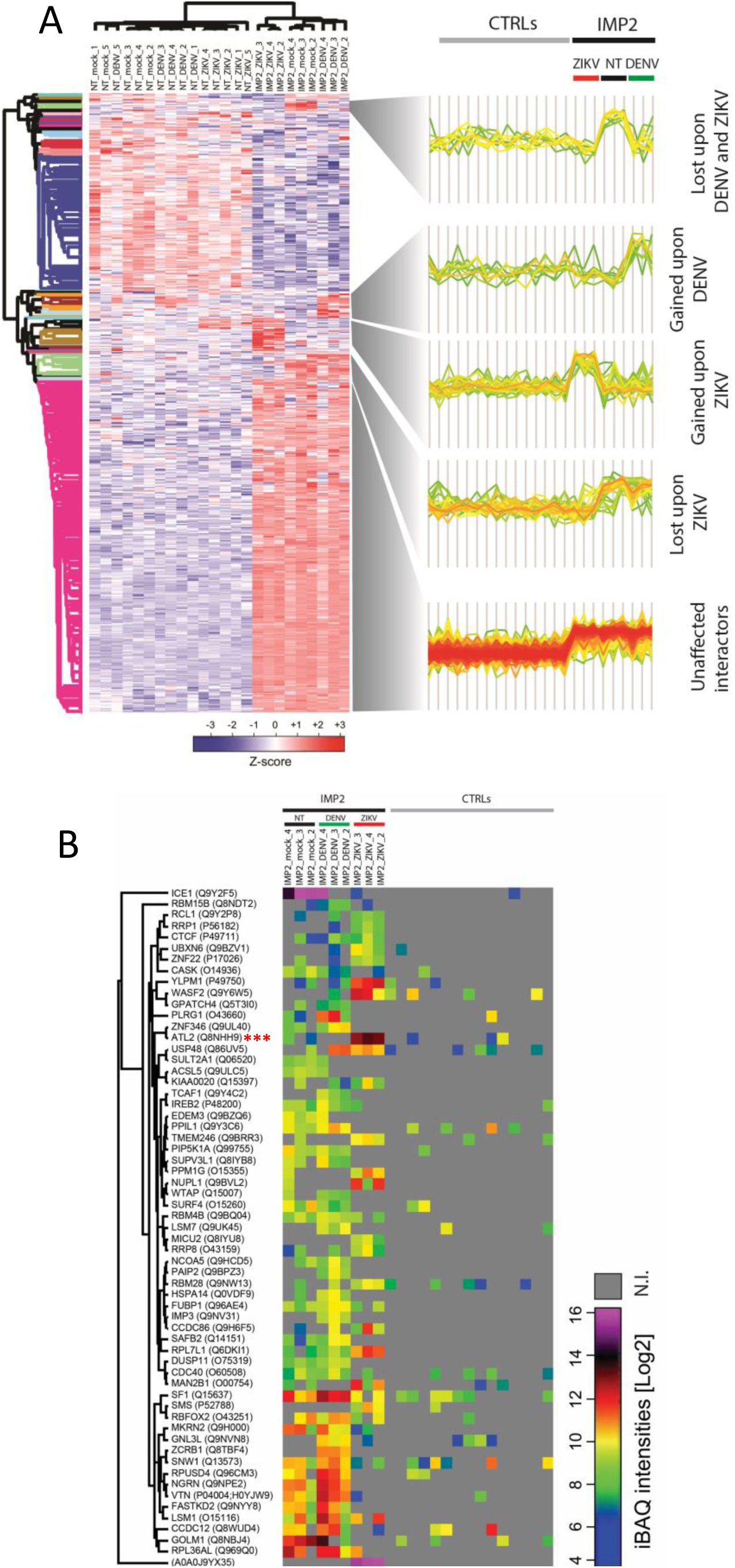
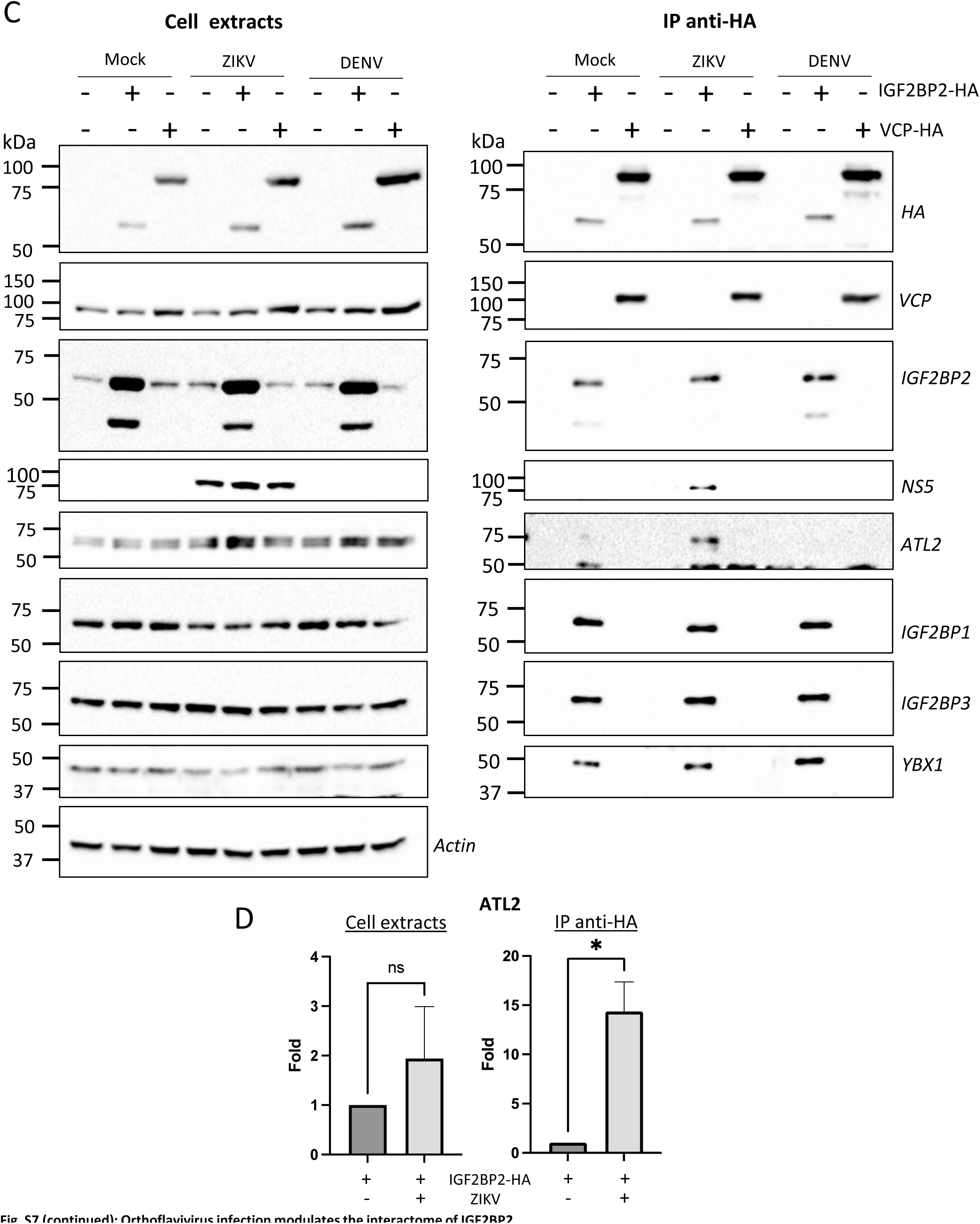
Orthoflavivirus infection modulates the interactome of IGF2BP2. (A) Heatmap of z-scored protein intensities of all significantly regulated interacting-proteins (ANOVA, FDR < 0.05) across stimuli (mock, DENV-infected, ZIKV-infected) and baits (NT, IGFBP2). (B) Absolute intensity-based quantification (iBAQs) of protein abundance of differential interacting proteins of IGF2BP2 upon DENV or ZIKV infection (N.I.=not identified). (C) Huh7.5 cells expressing IGF2BP2-HA and control cells were infected with ZIKV H/PF/2013 at a MOI of 10 or left uninfected. Two days later, cell extracts were prepared and subjected to anti-HA immunoprecipitations. The resulting complexes were analyzed by western blotting for their abundance in the indicated proteins. (D) ATL2 levels were quantified and means of protein signals (normalized to actin (extracts) and IGF2BP2 (IP)) ± SEM are shown. *: p<0.05; ns: not significant (unpaired t-test).

## Notes

### Competing Interest Statement

The authors have declared no competing interest.

### Summary of Updates

Figures 2, 4, 8, S1, S5, S6 and S7 revised Authors list updated

